# An interim exploratory biomarker analysis of a Phase 2 clinical trial to assess the impact of CT1812 in Alzheimer’s disease

**DOI:** 10.1101/2024.02.16.578765

**Authors:** BN Lizama, HA North, K Pandey, C Williams, D Duong, E Cho, V Di Caro, L Ping, K Blennow, H Zetterberg, J Lah, AI Levey, M Grundman, AO Caggiano, NT Seyfried, ME Hamby

## Abstract

CT1812 is a novel, brain penetrant small molecule modulator of the sigma-2 receptor (S2R) that is currently in clinical development for the treatment of Alzheimer’s disease (AD). Preclinical and early clinical data show that, through S2R, CT1812 selectively prevents and displaces binding of amyloid beta (Aβ) oligomers from neuronal synapses and improves cognitive function in animal models of AD. SHINE is an ongoing Phase 2 randomized, double-blind, placebo-controlled clinical trial (COG0201) in patients with mild to moderate AD, designed to assess the safety and efficacy of 6 months of CT1812 treatment. To elucidate the mechanism of action in AD patients and pharmacodynamic biomarkers of CT1812, the present study reports exploratory cerebrospinal fluid (CSF) biomarker data from an interim analysis of the first set of patients in SHINE (part A). Untargeted mass spectrometry-based discovery proteomics can detect more than 2,000 proteins in patient CSF and has documented utility in accelerating the identification of novel AD biomarkers reflective of diverse pathophysiologies beyond amyloid and tau and enabling identification of pharmacodynamic biomarkers in longitudinal interventional trials. We leveraged this technique to analyze CSF samples taken at baseline and after 6 months of CT1812 treatment. Proteome-wide protein levels were detected using tandem mass tag-mass spectrometry (TMT-MS), change from baseline was calculated for each participant, and differential abundance analysis by treatment group was performed. This analysis revealed a set of proteins significantly impacted by CT1812, including pathway engagement biomarkers (i.e., biomarkers tied to S2R biology) and disease modification biomarkers (i.e., biomarkers with altered levels in AD vs. healthy control CSF but normalized by CT1812, and biomarkers correlated with favorable trends in ADAS-Cog11 scores). Brain network mapping, Gene Ontology, and pathway analyses revealed an impact of CT1812 on synapses, lipoprotein and amyloid beta biology, and neuroinflammation. Collectively, the findings highlight the utility of this method in pharmacodynamic biomarker identification and providing mechanistic insights for CT1812, which may facilitate the clinical development of CT1812 and enable appropriate pre-specification of biomarkers in upcoming clinical trials of CT1812.

**HIGHLIGHTS:**

- Effects of CT1812 on AD patients were investigated in a randomized Phase 2 clinical trial
- Pharmacodynamic biomarkers of CT1812 were identified through unbiased analysis of proteomics quantitation data acquired using TMT Mass Spectrometry (TMT-MS)
- CT1812 normalized a set of biomarkers altered in AD
- Findings provide proof of mechanism that CT1812 impacts synapse, inflammation, amyloid-related processes

## INTRODUCTION

There is a huge unmet medical need to develop novel therapeutics for Alzheimer’s disease (AD), the most prevalent cause of dementia and neurodegeneration, given the limited treatment options available. A robust pipeline of investigational therapeutics in both preclinical and clinical development addresses a range of targets and biological processes (Cummings et al., 2023). Chief among the core pathologies targeted by therapeutics under development are amyloid beta (Aβ), tau, inflammatory processes, and neurotransmission. Significant advancements in the field have occurred recently, with full approval of lecanemab and accelerated approval of aducanumab, both Aβ-lowering immunotherapy approaches (Budd Haeberlein et al., 2022; van Dyck et al., 2023). Although these therapeutics lower Aβ and are able to slow disease progression by about 30%, patients continue to experience cognitive decline (van Dyck et al., 2023). Thus, there remains a need to further impact the disease course, and additional therapeutic approaches are imperative to pursue beyond these or in combination with existing options.

CT1812, a novel brain penetrant small molecule modulator of the sigma-2 receptor (S2R), is currently in clinical development for AD. In preclinical studies, CT1812 selectively prevented and displaced the binding of Aβ oligomers from their neuronal synaptic receptors, restored neuronal function in a trafficking assay, and rescued synapse loss in neuronal cultures (Izzo et al., 2021). In a mouse model of AD, CT1812 restored cognition to healthy control levels (Izzo et al., 2021). Early proof of mechanism of Aβ displacement by CT1812 has been demonstrated in a small single-dose clinical trial in AD patients (LaBarbera et al., 2023); however, assessment of longer treatment courses are needed to understand the impact of CT1812 over extended periods of time, and as CT1812 treatment progresses through early-phase clinical trials in humans, the biological impact of CT1812 still lacks a comprehensive understanding.

SHINE (COG0201; NCT03507790) is an ongoing Phase 2 randomized, double-blind, placebo-controlled clinical trial studying the safety, tolerability, and efficacy of 6 months treatment with CT1812 in patients with mild to moderate AD. An exploratory aim of SHINE is to analyze and identify cerebrospinal fluid (CSF)- based biomarkers to elucidate CT1812’s mechanism of action in AD patients and identify pharmacodynamic biomarkers of CT1812 efficacy. The present manuscript details the CSF biomarker findings from the first 24 participants in SHINE as an interim analysis (SHINE-A).

A rapidly expanding field, driven by research at both academic institutions and by the pharmaceutical/ biotechnology industry, strives to identify and assess biomarkers in AD patients to facilitate diagnosis and drug development. The development of new diagnostic biomarkers in recent years enabled the accelerated FDA approval of the first disease modifying therapy for AD, aducanumab (Aduhelm), which was largely based upon biomarker evidence of target engagement through the measurement of Aβ plaque levels, a hallmark of AD of which a reduction is thought to reflect disease modification (Cavazzioni, 2021).

While hypothesis-driven assessment of individual biomarkers with known links to AD has borne fruit, the identification of novel AD biomarkers through large-scale discovery proteomics has dramatically accelerated the field. Unbiased discovery proteomics, using tandem mass tag (TMT) mass spectrometry (MS), can measure levels of more than 2,000 proteins from a single CSF sample and can enable assessment of longitudinal changes in abundance across all proteins detected (Higginbotham et al., 2020). This technique enables the procurement of rich datasets from which novel biomarkers linked to several key pathophysiological processes perturbed in the disease (Higginbotham et al., 2020; Johnson et al., 2022) and modulated by interventional therapeutics can be discovered (Levey et al., 2022).

Here, we have utilized this approach in the context of the SHINE Phase 2 clinical trial to further elaborate the mechanism of action of CT1812 while simultaneously identifying pharmacodynamic biomarkers of CT1812 to support its clinical development. Utilizing TMT-MS proteomics, we identified and quantified more than 2,000 proteins in CSF collected from the first 24 participants of SHINE. To validate our approach, we first compared proteomic quantifications identified through TMT-MS against results from known, quantitative immunoassays, confirming strong correlations across platforms. Following this, we analyzed the distinct CSF proteins and pathways that were altered in patients treated with CT1812 and found them linked to brain-based, disease-related biologies such as synapses, immune function, and Aβ. Pearson correlation analyses revealed biomarkers correlated with slower decline in cognitive performance in drug-treated patients, as well as that correlated with changes in CSF levels of the AD hallmark biomarker Aβ. Together, the findings reported here speak to the primary biological processes that CT1812 may modify, providing insights into the mechanism by which CT1812 impacts AD.

## METHODS

### Clinical trial design

SHINE (COG0201) is an international, multi-center, randomized, double-blind, placebo-controlled parallel group 36-week Phase 2 clinical trial of CT1812 in adults with mild to moderate AD. SHINE is designed to enroll approximately 144 participants; men and women aged 50-85 years with mild to moderate AD (MMSE 18-26) were screened for eligibility. AD diagnoses were confirmed by amyloid PET imaging or by CSF biomarkers measured at the screening visit. Participants were randomized 1:1:1 to receive placebo, or 100 or 300 mg CT1812, orally, once daily for 6 months (Figure 1A).

**Figure 1.**
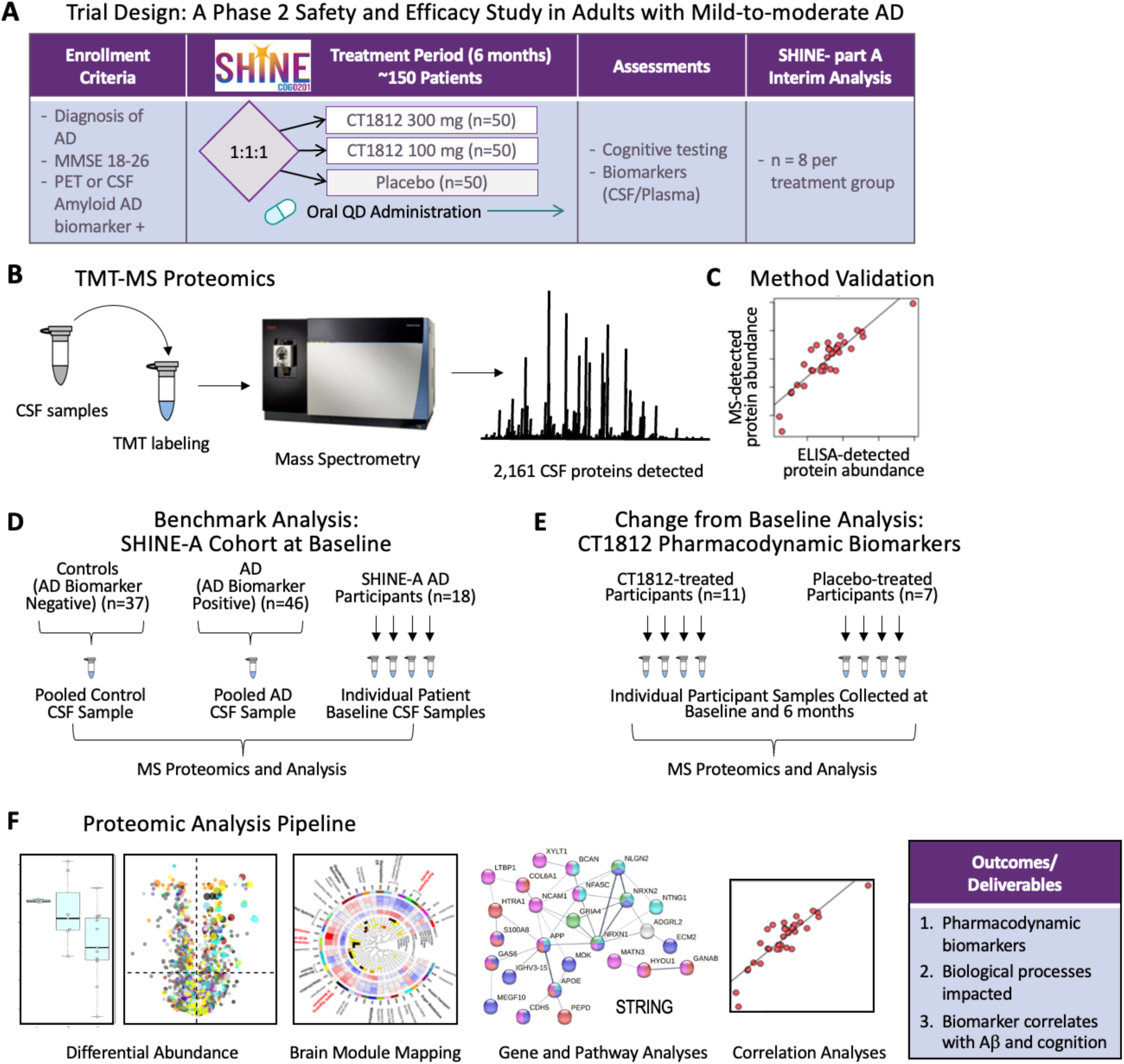
Study design. **A.** SHINE-A clinical trial design. **B.** Analysis of patient CSF samples using TMT-MS to measure >2,000 proteins. **C.** To validate the quantitative nature of the MS approach, proteins of interest were measured using clinically validated assays (ELISA) and correlation analyses were performed against the >2,000 proteins measured using TMT-MS. **D.** To benchmark the SHINE-A cohort at baseline against a known AD cohort and to further validate the quantitative use of TMT-MS, baseline SHINE-A CSF proteomes were compared to those of pooled Emory control (healthy) individuals and pooled Emory AD patient reference population CSF samples. **E.** To quantify changes in CSF proteome following treatment with CT1812 and identify possible CT1812 pharmacodynamic biomarkers, change from baseline CT1812-and placebo-treated patient CSF proteomes were compared. **F** Proteome analysis pipeline. Differentially abundant proteins were identified, mapped to established brain modules, assessed for interconnectivity and biological relevance, and using STRING, Metacore, and Gene Ontology. Proteins abundances that correlated with AD traits such as Aβ and ADAS-Cog11 were also identified and analyzed.

The primary and secondary objectives of this trial are to evaluate the safety and efficacy of CT1812. Exploratory outcome measures include assessment of CSF and plasma biomarker changes longitudinally. In addition to clinical assessments, biomarker assessments were performed on CSF samples taken by lumbar puncture at baseline and 6 months. Outcome measures included safety and tolerability, cognitive function as measured by the Alzheimer’s Disease Assessment Scale – Cognitive Subscale 11-item version (ADAS-Cog11), and CSF biomarkers of disease modification.

The present manuscript reports the interim analysis of the CSF TMT-MS discovery proteomics data gathered from the first 24 participants evaluated as part A of the study. Biomarker analyses were performed only on samples from participants for whom timepoint-matched biofluids were available (i.e., of 24 total participants, 18 CSF samples were successfully collected at both baseline and 6 months).

### Safety

CT1812 has been found to be generally safe and well tolerated in clinical trials: CT1812 was found to be safe and well tolerated in a Phase 1 placebo-controlled SAD/MAD safety study in healthy volunteers (NCT03716427) (Grundman et al., 2019) and in a placebo-controlled Phase 1a/2 trial in subjects with mild to moderate AD (NCT02907567). CT1812 has also recently been evaluated in a 6-12 month Ph1/2 randomized, double-blind, placebo-controlled parallel-group trial in subjects with mild to moderate AD (NCT03493282) (van Dyck et al., 2024).

Safety and tolerability are the primary outcome measures in the SHINE trial. Although not the focus of this manuscript, safety data were collected for the SHINE-A interim analysis to add to CT1812 safety data that has been previously collected. In this interim analysis of findings from the first 24 participants in the SHINE trial, one subject withdrew from the study due to treatment-emergent adverse events. There were no serious adverse events attributed to study medication.

### Exploratory biomarker analyses

All biomarker analyses carried out for this manuscript were exploratory and for the purpose of identifying pharmacodynamic changes of CT1812, and only participants who were actively taking their treatment, as indicated by bioanalysis of drug exposure levels (herein referred to as treatment-compliant participants), were included in the analysis. For AD core biomarkers (neurogranin (Nrgn), synaptotagmin (Syt1), neurofilament light (NfL), and tau), mean and standard error of the mean (SEM) are shown, with Student’s t-test used to determine significance. TMT-MS differential abundance was assessed via ANOVA following determination of the log2 change from baseline.

### CSF Tandem-mass tag mass spectrometry (TMT-MS) proteomics measurements

CSF samples from SHINE-A participants for whom both baseline and end of study samples were collected were processed and analyzed using TMT-MS followed by unbiased quantification (N = 18 participants: 7 = placebo, 4 = 100 mg CT1812, 7 = 300 mg CT1812) as previously described (Modeste et al., 2023) (Figure 1B). Using TMT-MS, 2,161 proteins were detected reliably across all SHINE-A CSF samples using the <50% detection exclusion criterion.

Protein digestion of CSF: First, 70 μl of CSF was digested with lysyl endopeptidase (LysC) and trypsin, reduced and alkylated with 1.4 μl of 0.5 M tris2(-carboxyethyl)-phosphine (Thermo Fisher) and 7 μl of 0.4 M chloroacetamide in a 90 °C water bath for 10 min, following with bath sonication for 5 min. Aoer lepng samples cool to room temperature, 78 μl of 8 M urea buffer (8 M urea, 10 mM Tris, 100 mM NaH2PO4, pH 8.5) and 3.5 μg of LysC (Wako) were added to each sample, resulting in a final urea concentration of 4 M. The samples were mixed, spun down and incubated overnight at 25 °C for digestion with LysC. The following day, samples were diluted to 1 M urea with solution containing 468 μl of 50 mM ammonium bicarbonate and 7 μg of Trypsin (Thermo Fisher). The samples were subsequently incubated overnight at 25 °C for digestion with trypsin, acidified to a final concentration of 1% formic acid and 0.1% trifuoroacelc acid. This was immediately followed by desalting on 30 mg HLB columns (Waters) and then eluted with 1 ml of 50% acetonitrile (I). To normalize protein quantification across batches, 100 μl was taken from all CSF samples and then combined to generate a pooled sample. This pooled sample was then divided into global internal standards (GIS). All individual samples and the pooled standards were then dried using a speed vacuum (Labconco).

TMT labeling of CSF peptides: All CSF samples were balanced for treatment group, sex, and APOE status across 4 batches and loaded on to a 16-plex TMT (TMTpro) kit (Catalog# A44520 and Lot # UI292951). Care was taken to ensure baseline and 6-month CSF samples from same participant were included in the same batch. Further, one GIS pool and AD and Control standard pools created by aggregating CSF from age matched AD and Control cohorts from Emory Goizueta Alzheimer’s Disease Research Center (ADRC) was added per batch. In preparation for labeling, each CSF peptide digest was resuspended in 75 μl of 100 mM triethylammonium bicarbonate (TEAB) buffer. Meanwhile, 5 mg of TMT reagent was dissolved into 200 μl of I. Once both were in suspension, 15 μl of TMT reagent solution was subsequently added to the resuspended CSF peptide digest. Aoer 1 h, the reaction was quenched with 4 μl of 5% hydroxylamine (Termo Fisher Scientific, 90,115) for 15 min. The peptide solutions were then combined according to the batch arrangement. Finally, each TMT batch was desalted with 60 mg HLB columns (Waters) and dried via speed vacuum (Labconco).

High-pH peptide fractionation: Dried samples were re-suspended in high pH loading buffer (0.07% vol/vol NH4OH, 0.045% vol/vol FA, 2% vol/vol I) and loaded onto a Water’s BEH column (2.1 mm×150 mm with 1.7 µm particles). A Vanquish UPLC system (Thermo Fisher Scientific) was used to carry out the fractionation. Solvent A consisted of 0.0175% (vol/vol) NH4OH, 0.01125% (vol/vol) FA, and 2% (vol/vol) I; solvent B consisted of 0.0175% (vol/ vol) NH4OH, 0.01125% (vol/vol) FA, and 90% (vol/vol) I. The sample elution was performed over a 25 min gradient with a flow rate of 0.6 mL/min with a gradient from 0 to 50% solvent B. A total of 192 individual equal volume fractions were collected across the gradient. Fractions were concatenated to 96 fractions and dried to completeness using vacuum centrifugation.

Mass spectrometry analysis and data acquisition: All samples (∼1 µg for each fraction) were loaded and eluted by 1200Ultimate U3000 RSLCnano (Thermo Fischer Scientific) with an in-house packed 15 cm, 150 μm i.d. capillary column with 1.7 μm CSH (Water’s) over a 35 min gradient. MS was performed with a high-field asymmetric waveform ion mobility spectrometry (FAIMS) Pro front-end equipped Orbitrap Eclipse (Thermo Fisher) in positive ion mode using data-dependent acquisition with 1 s top speed cycles for each FAIMS compensative voltage. Each cycle consisted of one full MS scan followed by as many MS/MS events that could o within the given 1 s cycle time limit. MS scans were collected at a resolution of 120,000 (410– 1600 m/z range, 4×10^5 AGC, 50 ms maximum ion injection lme, FAIMS compensative voltage of -45 and -65). Only precursors with charge states between 2+ and 6+ were selected for MS/MS. All higher energy collision-induced dissociation (HCD) MS/MS spectra were acquired at a resolution of 30,000 (0.7 m/z isolation width, 35% collision energy, 1.25×10^5 AGC target, 54 ms maximum ion time). Dynamic exclusion was set to exclude previously sequenced peaks for 20 s within a 10-ppm isolation window.

Database search and protein quantification: All raw files were analyzed using the Proteome Discoverer Suite (v.2.4.1.15, Thermo Fisher). MS/MS spectra were searched against the UniProtKB human proteome database (downloaded in 2015 with 90,300 total sequences). The Sequest HT search engine was used to search the RAW files, with search parameters specified as follows: fully tryptic specificity, maximum of two missed cleavages, minimum peptide length of six, fixed modifications for TMTPro tags on lysine residues and peptide N-termini (+304.304.2071 Da) and carbamidomethylation of cysteine residues (+57.02146 Da), variable modifications for oxidation of methionine residues (+15.99492 Da), serine, threonine and tyrosine phosphorylation (+79.966 Da) and deamidation of asparagine and glutamine (+0.984 Da), precursor mass tolerance of 10 ppm and a fragment mass tolerance of 0.05 Da. Percolator was used to filter peptide spectral matches and peptides to an FDR<1%. Following spectral assignment, peptides were assembled into proteins and were further filtered based on the combined probabilities of their constituent peptides to a final false discovery rate (FDR) of 1%. Peptides were grouped into proteins following strict parsimony principle.

### Validation of TMT-MS as a quantitative approach

Key AD core proteins, related to AD pathology, were measured from participant CSF samples using clinically validated assays (Blennow & Zetterberg, 2018). NfL was measured using Uman Diagnostics; t-Tau, Nrgn, and Syt1 were measured using Lumipulse (herein referred to as qNfl, qTau, qNrgn, qSyt1).

Correlation analyses were performed for each of the 2,161 proteins reliably detected across ≥ 50% samples using TMT-MS (Figure 1B-C). The most strongly correlated proteins (e.g., r ≥ |0.50| or |0.70|, p ≤ 0.01 or ≤ 0.05) for each quantitative approach were analyzed using STRING as indicated in each figure legend.

### Benchmarking of proteomes from SHINE-A patients to AD reference population

To determine whether the SHINE-A AD participant population CSF proteome resembled those from a previously well-characterized AD cohort (Watson et al., 2023), proteomes were compared, within-study, to pooled AD biomarker-positive (AD) and age-matched, non-demented biomarker-negative (control) CSF reference standards from the ADRC (Figure 1D). Briefly, AD and control CSF reference standards were generated based on participant Aβ42, total tau (tTau), and pTau181 levels measured using the Elecys platform (Watson et al., 2023). Equal volumes of CSF were combined from each participant (approximately 45 individual samples were combined to form each pooled sample), resulting in a final sample of approximately 50 ml per pool before aliquoting. The AD reference standard pool had low levels (average) of Aβ42 (482.6 pg/mL) and high total tau (341.3 pg/mL) and pTau181 (33.1 pg/mL), and the control CSF reference standard pool had relatively high levels of Aβ42 (1457.3 pg/mL) and low total tau (172.0 pg/mL) and pTau181 (15.1 pg/mL) (Watson et al., 2023). These reference samples were analyzed using TMT-MS in the same manner and on the same plexes as SHINE-A participant CSF and the abundances of AD-related proteins were compared (Figure 3C).

Additionally, proteins that were found to be differentially expressed between AD and control reference standard CSF proteomes were subjected to pathway analysis and brain mapping, as described below.

### Proteomic Analysis Pipeline

#### Identification of pharmacodynamic biomarkers of CT1812

To identify pharmacodynamic biomarkers of CT1812, change from baseline CSF levels of CT1812- and placebo-treated proteins were compared for statistical significance via ANOVA and differentially expressed proteins (p ≤ 0.05) were identified.

#### Brain mapping

The identified differentially expressed proteins were mapped to one of the 44 modules associated with AD biology as described in a previously-published protein co-expression network that was built using proteomic data from brain samples of healthy individuals, asymptomatic and symptomatic AD cases (Johnson et al., 2022), using previously described methods (Guo Q et al., 2023) (Figure 1F).

#### Pathway analyses

Proteins meeting the statistical threshold for each comparison were assessed using STRING (version 11.5 or 12.0, indicated for each figure) and Metacore (version 23.3.71400) (Figure 1F). All proteins without names (e.g., “0”) were excluded from the pathway analyses. The protein-protein interaction (PPI) network for each condition was exported and all pathway terms (e.g., for KEGG, GO Biological Processes, Reactome, etc.) were ranked according to strength or FDR for interpretation.

For visualization of the PPI network, low, medium, and high confidence was used as appropriate (as indicated for each figure). Unconnected nodes were excluded from the PPI maps.

### Pre-specified ADAS-Cog 11 statistical analyses

Statistical analysis of the cognitive assessment scale ADAS-Cog11 data was performed according to the pre-specified statistical analysis plan at the time (V1). The participants included in the exploratory efficacy assessment were the Modified Intent-To-Treat Population (mITT), which included all randomized subjects who receive any amount of study drug and who have a baseline and at least one post-baseline assessment of the ADAS-Cog 11 total score was used for the analysis. For the interim analysis, an ANCOVA model at each visit was run, with baseline ADAS-11 score in place of the composite baseline score. For the interim analysis, an analysis of covariance (ANCOVA) model at each visit was run. This model included treatment as the main effect, baseline z-score of the corresponding composite scores, and APOE ε4 (+ or −) status at baseline. For endpoints where LOCF was used, the analyses were repeated using only observed data. The primary analysis of the ADAS-Cog compared the two active arms combined to the placebo arm at day 182, using a two-sided test at the alpha=0.05 level of significance

## RESULTS

### Construct validation of TMT-MS as a quantitative measure using CSF samples

Using TMT-MS, 2,161 proteins were detected reliably across all SHINE-A CSF samples using the < 50% detection exclusion criterion. To validate the method as quantitative, Pearson correlation analyses were performed across all samples between levels for each of these 2,161 CSF proteins with levels of the core AD biomarkers total tau (tTau; qTau [quantitative tau]), neurofilament light (NfL; qNfL), synaptotagmin 1 (Syt1; qSyt1), and neurogranin (Nrgn, qNrgn) (Blennow & Zetterberg, 2018), as measured using independent, clinically validated, quantitative methods (Blennow & Zetterberg, 2019). Notably, a high congruency was observed between the levels detected for each core biomarker using the two methods, as depicted via the Pearson correlation coefficient (r) heatmap (SYT1, r = 0.69; NRGN, r = 0.90; NEFL, r = 0.88; MAPT, r = 0.85; p ≤ 0.001) (Figure 2A): Scatterplots of correlations of each individual sample for validated assay-measured and corresponding MS-measured proteins are shown (Figure 2B). For each of these core AD biomarkers, many additional MS-detected proteins were found to be highly and significantly correlated (r ≥ |0.50| and p ≤ 0.05), both positively and negatively (Figure 2C).

**Figure 2.**
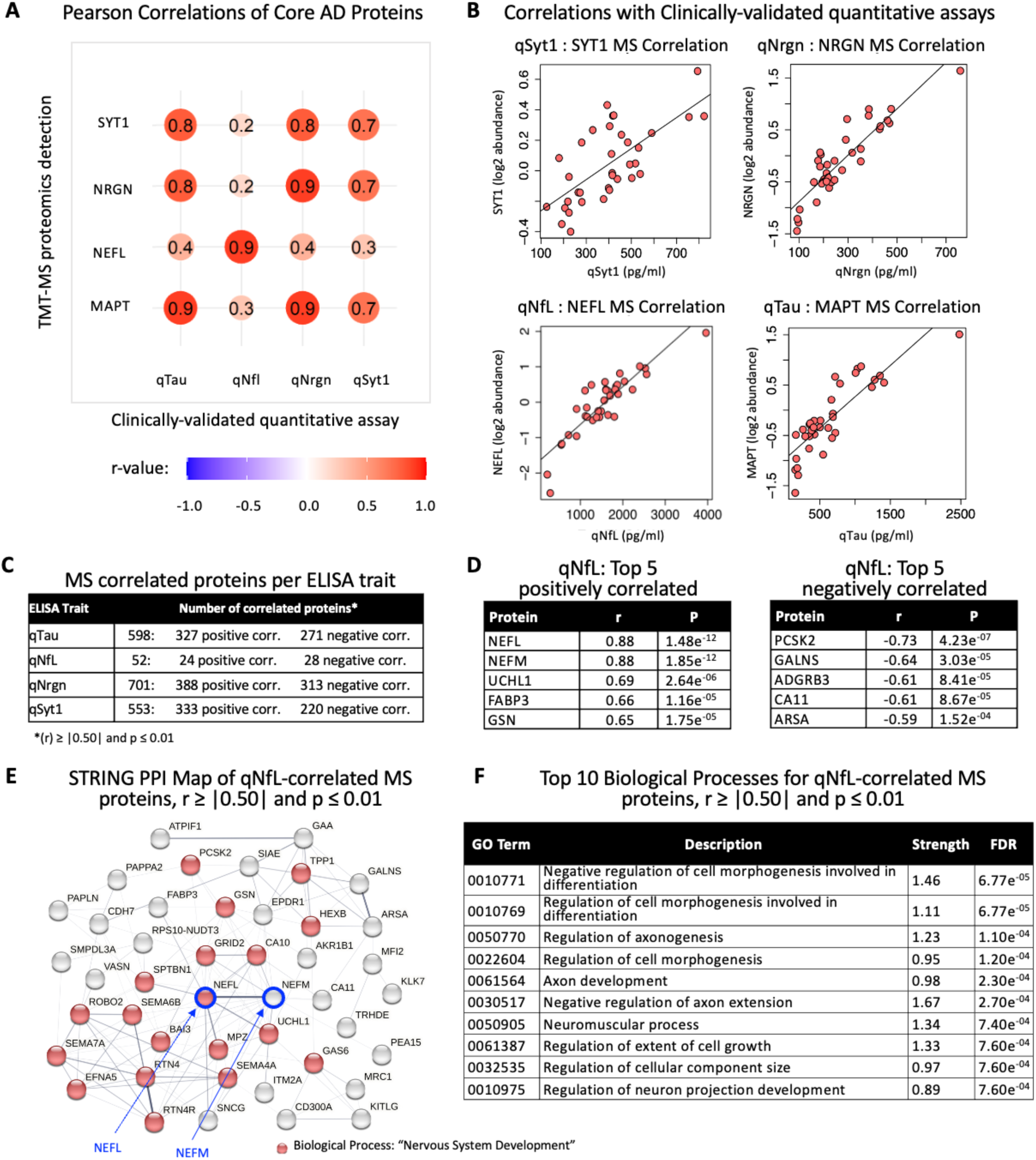
Validation of TMT-MS proteomics as a quantitative method. Pearson correlation analysis compared levels of AD core biomarkers in SHINE-A patient CSF samples measured using clinically validated assays (Blennow et al., 2019) to those measured using TMT-MS proteomics (N=36: protein levels in 2 samples, baseline and 6 months, per participant). **A**. The strength of correlations between levels of key proteins related to AD from each method are plotted on a heat map to visualize the high degree of correlation between ELISA and quantitative TMT-MS. **B.** Scatter plots illustrate the congruency between levels of each core AD biomarker examined (quantitative (q) NfL (qNfL), qTau, qNrgn, qSyt1), and the levels detected via TMT-MS. **C.** For each AD core biomarker measured using a clinically validated assay, the number of TMT-MS proteins highly correlated (r ≥ |0.50| and p ≤ 0.01), along with directionality (either positively (+) or negatively (-) correlated) is indicated. **D**. The top 5 strongest positively and negatively correlated TMT-MS proteins to qNfL are listed with NEFL and NEFM identified as the most strongly correlated. **E.** The STRING (v11.5) protein-protein interaction (PPI) map of all significantly correlated proteins (n = 52; r ≥ |0.50| and p ≤ 0.01) is depicted; NEFL and NEFM are central hubs. **F.** Gene Ontology analysis of these 52 proteins identified the top-most enriched 10 pathways (ranked by FDR p-value); processes related to axon dynamics are strongly represented.

Characteristics of these correlates were analyzed for each of these core AD biomarkers. Figure 2D-F demonstrates the analysis performed for NfL, a biomarker of neurodegeneration that is increased in AD patient CSF (Zetterberg et al., 2016) (Figure 3A). Of the 52 TMT-MS proteins robustly correlated to qNfL assessed via ELISA (r ≥ |0.50| and p ≤ 0.05), the top 5 positive and negative correlates, with strong significance, are shown (Figure 2D); NEFL (r = 0.88, p = 1.48e^-12^), NEFM (r = 0.88, p = 1.85e^-12^), PCSK2 (r = - 0.73, p = 4.23e^-07^, UCHL1 (r = 0.69, p = 2.64e^-06^) and FABP3 (r = 0.66, p = 1.16e^-05^) are the topmost correlated proteins. Scatter plots illustrate the relationship across individual CSF samples and depict the strong correlation of qNfL to NEFL (Figure 2B, bottom left) and NEFM (Supplemental Figure 1A). The 52 correlates were analyzed using STRING to illustrate protein-protein interactions (Figure 2E-F), which were found to show interactions among the most correlated proteins and to include processes concordant with the cellular localization and function of the axonally expressed NfL (Laser-Azogui et al., 2015): “Regulation of axonogenesis” (ranked 3^rd^), “Axon development” (ranked 5^th^), and “Negative regulation of axon extension” (ranked 6^th^).

**Figure 3.**
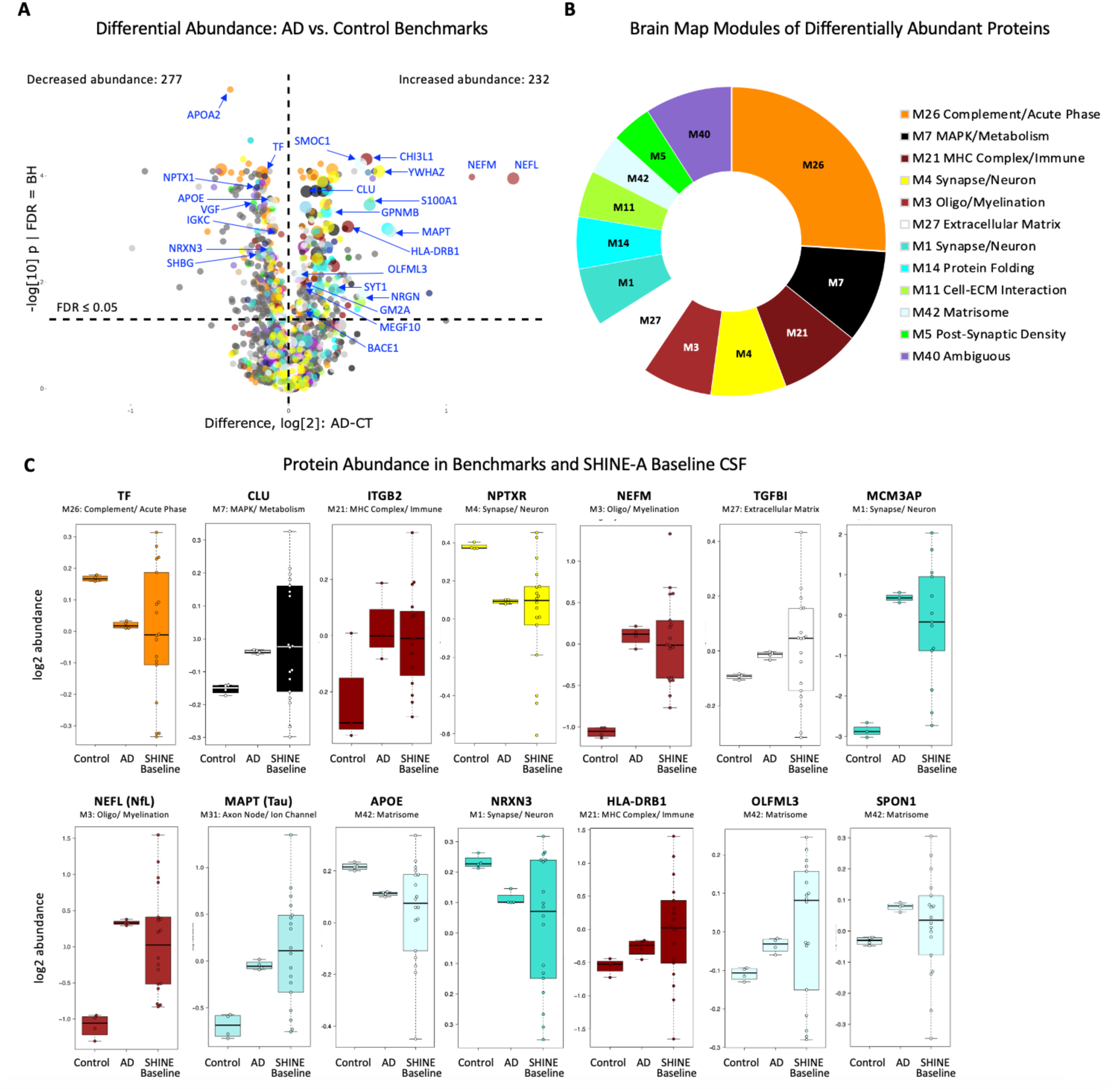
Identification of proteins disrupted in AD, and characterization of the SHINE-A proteomic phenotype at baseline. **A.** Differential abundance of proteomes as measured vis TMT-MS from pooled AD biomarker positive CSF compared to pooled control (healthy) biomarker negative CSF was assessed (AD vs. control; FDR ≤ 0.05, BH = Benjamini-Hochberg correction). Log2 abundance and significance for each protein detected is shown on the volcano plot (each colored dot corresponds to a protein; colors correspond to module assignment colors in (B); larger circles indicate AD priority biomarkers. 509 proteins (277 decreased and 232 increased) were found to be significantly differentially abundant. Proteins of interest are indicated with royal blue arrows: APOA2, apolipoprotein A2; APOE, apolipoprotein E; BACE1, Beta-Secretase 1; CHI3L1, chitinase-3-like protein 1; CLU, clusterin; GM2A, ganglioside activator; GPNMB, Glycoprotein nonmetastatic melanoma protein B; HLA-DRB1, human leukocyte antigen class II histocompatibility D related beta chain; IGKC, Immunoglobulin kappa constant; MAPT, microtubule-associated protein tau; MEGF10, Multiple EGF-like-domains 10; NEFL, neurofilament light chain; NEFM, neurofilament medium chain; NRGN, neurogranin; NRXN3, neurexin 3; OLFML3, olfactomedin-like protein 3; TF, transferrin; S100A1, S100 calcium binding protein A1; SHBG, sex hormone binding globulin; SMOC1, SPARC-related modular calcium-binding protein 1; SYT1, synaptotagmin 1; YWHAZ, 14-3-3 protein zeta/delta; VGF, VGF nerve growth factor inducible; NPTX1, neuronal pentraxin 1). **B.** The proteins identified as significantly differentially abundant (FDR ≤ 0.01) were mapped to previously established AD co-expression modules (Johnson et al., 2022), and the top related categories to which the proteins belong are shown on the pie chart. The greatest representation of proteins ascribed to the module is reflected by the relative size of the slice of pie, and the list ordered from top to least most represented. **C.** Abundances of AD-related genes were compared between healthy control, the Emory AD and control reference standards and the SHINE-A baseline CSF proteomes to determine how SHINE-A participant CSF compares to CSF of either healthy or AD known populations. Data is presented as mean abundance (line within box) +/-SEM (box height) and +/-SD (error bars). Proteins shown: Complement/ Acute phase protein TF (orange); MAPK/ Metabolism protein CLU (black); MHC Complex/ Immune protein ITGB2 (beta 2 integrin; maroon); Synapse/ Neuron protein NPTXR (neuronal pentraxin receptor; yellow); Oligo/ Myelination protein NEFM (light maroon); Extracellular Matrix protein TGFB1 (transforming growth factor-beta-induced; white); Synapse/ Neuron protein MCM3AP (minichromosome maintenance complex component 3 associated protein, aqua), Oligo/ Myelination protein NEFL (light maroon); Axon Node/ Ion Channel protein MAPT (light aqua); Matrisome protein APOE (light cyan); Synapse/ Neuron protein NRXN3 (turquoise); MHC Complex/ Immune protein HLA-DRB1 (maroon); and Matrisome protein OLFML3 (light cyan).

As for qNfl, STRING Pathway analyses of the set of proteins highly correlated (r ≥ |0.50| and p ≤ 0.01) to the other core AD biomarkers Syt1, Nrgn, and Tau were performed to identify protein-protein interactions and networks. The topmost significant Biological Process Gene Ontology (GO) and Molecular Function GO terms for each network of correlates are displayed (Supplemental Figure 1B).

The STRING protein-protein interaction map for the post-synaptic protein involved in memory, neurogranin (Nrgn), displays the most inter-connected proteins out of the 166 TMT-MS proteins found be significantly correlated with qNrgn using a criterion of r ≥ |0.70| and p ≤ 0.01 (Supplemental Figure 1C).

Nrgn itself is a highly interconnected node as indicated via the multiple of connecting lines within this pathway map of correlates (Supplemental Figure 1C). In this network, more than half of correlates (79 of 155 nodes on map) were found to be related to the GO molecular function “protein binding” (blue nodes), consistent with Nrgn being a calmodulin-binding protein (Prichard et al., 1999), and 29 of 155 nodes on map were related to the GO cellular component “synapse” (red nodes), as expected for its postsynaptic localization (Chang et al., 1997). An expanded list of the top-ranked Cellular Component terms is shown (Supplemental Figure 1D). Nrgn correlates were found to be highly represented within “vesicle,” “extracellular exosome,” or similar compartments, again, as expected given Nrgn is found in neuronally-derived exosomes in neurodegenerative conditions (Badhwar & Haqqani, 2020; Goetzl et al., 2016; Winston et al., 2016, 2018), and with GO terms “Neuron to neuron synapse,” “Postsynapse,” or similar compartments, also as expected for this postsynaptic protein. The Cellular Component terms associated with the TMT-MS correlates for both qSyt1 (Supplemental Figure 1E) and qTau (Supplemental Figure 1F) were similarly related to the known functions of these AD biomarker proteins – e.g., synaptic and axonal-related pathways, respectively (Y. Park & Ryu, 2018; Pooler et al., 2014). Altogether, these analyses support the utility of this method in its ability to provide quantitative results from which rich biological insights can be gleaned.

### Identification of disease-associated proteins and characterization of SHINE-A cohort baseline CSF proteome

To understand whether the SHINE-A AD cohort was prototypical of AD at baseline, we first analyzed, within the same TMT-MS run, the proteomes of the well-characterized pools of CSF from biomarker validated cases - either positive (AD) or negative (control) for canonical AD biomarkers - collected at the Emory Goizueta ADRC, described above (Watson et al., 2023), as benchmark reference standards (Figure 1D; Figure 3). Differential abundance analysis of TMT-MS-measured proteins comparing pooled AD vs. the pooled control CSF replicated earlier reports (Higginbotham et al., 2020). For example, proteins disrupted in AD CSF pool relative to control pool included AD biomarkers such as MAPT (tTau) and NEFL (NfL), which can be visualized on the volcano plot (Figure 3A). Brain mapping was performed to understand how the CSF proteins align with co-expression modules defined by protein co-expression as previously established in brain AD proteomics studies (Johnson et al., 2022). Integration of CSF and brain proteomes revealed that differentially abundant proteins in AD CSF primarily overlap in AD co-expression modules in brain (Figure 3B), with the top represented modules being “Complement/ Acute phase inflammatory” (M26, orange), “MAPK/ Metabolism” (M7, black), “MHC Complex/ Immune” (M21, maroon), “Synapse/ Neuron” (M4, yellow), “Oligo/ Myelination” (M3, red), and “Extracellular matrix” (M27, white).

For each of the top represented AD modules (Figure 3B), levels of representative biomarkers were plotted in the reference benchmark CSF across technical replicates in each batch (control and AD) to highlight the technical coefficient of variation and difference in effect size. We also assess within-CSF protein differences across the SHINE-A participants at baseline prior to treatment (Figure 3C) to assess similarity or dissimilarity to the AD or control pools. For the most part, the means of the SHINE-A baseline CSF biomarker abundances were found to be congruent with those from the benchmark AD biomarker positive pool and distinct from the healthy control AD biomarker negative pool for a number of proteins typically dysregulated in AD, including core AD biomarkers NEFL and MAPT (Tau), suggesting that the SHINE-A participant cohort is typical of patients with AD. For example, CLU (clusterin [apolipoprotein J/ APOJ], black), neuronal pentraxin receptor (NPTXR, yellow), NEFM and NEFL (maroon), and MAPT (tau, light aqua) are AD-related proteins that were found to be expressed similarly in the AD control population and the SHINE-A baseline population relative to the healthy control population. Notably, the SHINE-A participant population was heterogeneous at baseline (Figure 3B, see range of protein levels in box plot), as expected, which may reflect natural biological variation across patients and/or may reflect the presence of AD subpopulations within the cohort (Tijms et al., 2024).

### Identification and mapping of pharmacodynamic biomarkers of CT1812

To ascertain the pharmacodynamic effect of CT1812 on AD CSF proteomes, following TMT-MS, the fold change differences from baseline (pre-vs. post-CT1812 treatment or placebo) were calculated per participant followed by differential abundance analysis (CT1812 vs. placebo; p ≤ 0.05) (Figure 4). A volcano plot displays 122 differentially abundant biomarkers significant in CT1812 vs. placebo, with 52 CSF proteins increased and 70 proteins decreased (Figure 4A). Notable proteins of interest that relate to AD or the CT1812 mechanism of action (see discussion) include fatty acid binding protein 1 (FABP1), and the intracellular cholesterol transporter NPC1 (higher in CT1812 vs. placebo CSF), as well as neurexin 2 (NRXN2), amyloid precursor protein (APP), and CLU (lower in CT1812 vs. placebo CSF) (Figure 4A).

**Figure 4.**
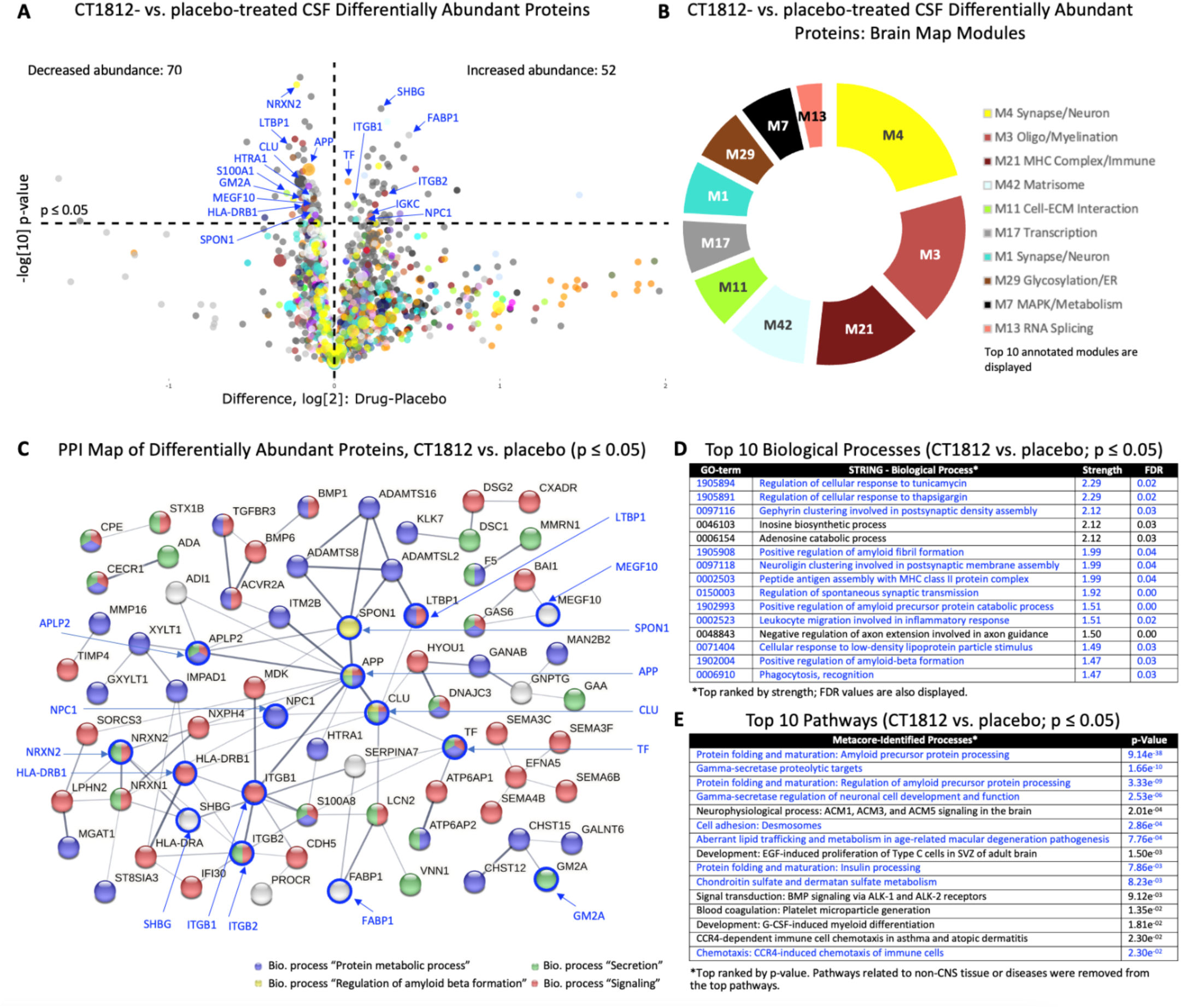
Analysis of differentially abundant proteins in CT1812- and placebo-treated patient CSF proteomes. **A.** Differential abundance of change from baseline levels of protein measured via TMT-MS was assessed via ANOVA (CT1812 vs. placebo; p≤0.05). Log2 abundance and significance for each protein detected is shown on the volcano plot (each colored dot corresponds to a protein; colors correspond to module assignment colors in (B); larger circles indicate AD priority biomarkers. 122 proteins (70 down, 52 up), were found to be significantly differentially abundant (CT1812 vs. placebo; p≤0.05). Proteins of interest are indicated with royal blue arrows: APP, amyloid precursor protein; CLU, clusterin; FABP1, fatty acid-binding protein 1; GM2A, ganglioside activator; HLA-DRB1, human leukocyte antigen class II histocompatibility D related beta chain; HTRA1, HtrA serine peptidase 1; IGKC, Immunoglobulin kappa constant; ITGB1, beta 1 integrin; ITGB2, beta 2 integrin; LTBP1, Latent-transforming growth factor beta-binding protein 1; MEGF10, multiple EGF-like-domains 10; NPC1, Niemann-Pick disease, type C1; NRXN2, neurexin 2; S100A1, S100 calcium binding protein A1; SHBG, sex hormone binding globulin; SPON1, spondin 1; TF, transferrin. **B**. The significantly (p ≤ 0.05) differentially abundant proteins were mapped to previously established AD co-expression modules (Johnson et al., 2022) and the related categories are shown on the pie chart. The greatest representation of proteins ascribed to the module is reflected by the relative size of the slice of pie, and the list ordered from top to least most represented **C**. STRING (v11.5) Protein-Protein Interaction (PPI) map of the significantly (p ≤ 0.05) differentially abundant proteins (medium confidence; proteins not connected not visualized). APP is a hub protein that was significantly lower in CT1812-treated patient CSF than in placebo-treated patient CSF. Other proteins of interest are indicated with royal blue arrows. Pathway analyses using both STRING (**D**) and Metacore (**E**) mapping show synapse, inflammatory, and amyloid pathways to be most significantly altered with CT1812 treatment when compared to placebo-treated controls.

Brain mapping of these proteins to AD co-expression modules demonstrated that proteins altered with CT1812 treatment vs. placebo (Figure 4B) map to disease-relevant brain modules that are highly represented in AD (Figure 3B; top 12 modules of 35 are shown). For example, many of the proteins changing in response to CT1812 treatment were highly represented in the “Synapse/ Neuron” module (Rank 1), followed by the “Oligodendrocytes/ Myelination” module (Rank 2), and the “MHC Complex/ Immune function” module (MHC, major histocompatibility complex; Rank 3).

To better understand the networks and pathways affected by CT1812, bioinformatic assessments were performed on significantly differentially abundant proteins (i.e., CT1812 vs. placebo; p ≤ 0.05) using STRING (Szklarczyk et al., 2023). The protein-protein interaction map, or network, (Figure 4C) displays the interconnectivity and relationships of these differentially abundant proteins to one another, with key functional roles of these proteins / pathways impacted highlighted (Figure 4D). Amyloid precursor protein (APP), a protein significantly decreased in abundance in CT1812-treated vs. placebo-treated CSF (Figure 4A), is a major hub protein in the protein-protein interaction map (Figure 4C). Top ranked Biological Processes GO terms were highly representative of endoplasmic reticulum/ protein folding, amyloid biology, inflammation, and synapse function (Figure 4D). Significantly altered proteins (p ≤ 0.05) were also assessed using Metacore pathway analysis to further verify the associations revealed by STRING, and to elucidate related functions and processes using a distinct, curated database of pathway function (Figure 4E).

Metacore analysis revealed pathways that were most significantly altered with CT1812 treatment to be amyloid biology-related (“Protein folding and maturation: APP processing,” “Gamma-secretase proteolytic targets,” “Gamma-secretase regulation of neuronal cell development and function,” and “Aberrant lipid trafficking and metabolism in age-related macular degeneration pathogenesis”), lending additional credence to CT1812’s defined mechanism as impacting trafficking and amyloid biology.

### Identification of biomarkers altered in AD that are normalized by treatment with CT1812

Leveraging the internal Emory AD and control CSF pooled reference standards within the TMT-MS proteomics study, we asked whether proteins that were increased or decreased in the AD pool (Figure 3A) were significantly affected by CT1812 treatment (Figure 4A) were normalized or shifted towards levels of healthy non-demented control pool by CT1812 treatment (Figure 4a). Twenty-one proteins were found to be significantly (p ≤ 0.05) normalized towards control levels in CT1812-vs. placebo-treated CSF (Figure 5A). Box plots (Figure 5B, C) illustrate two example proteins that were significantly increased in AD compared to control CSF (Figure 3), but significantly decreased in abundance in CT1812-treated vs. placebo-treated SHINE-A subject CSF (Figure 4). These included clusterin, CLU (Figure 5B) and HLA-DRB1 (Figure 5C), notably, both proteins that have been genetically linked to AD in large-scale genome-wide association studies (GWAS) (Lu et al., 2017).

**Figure 5.**
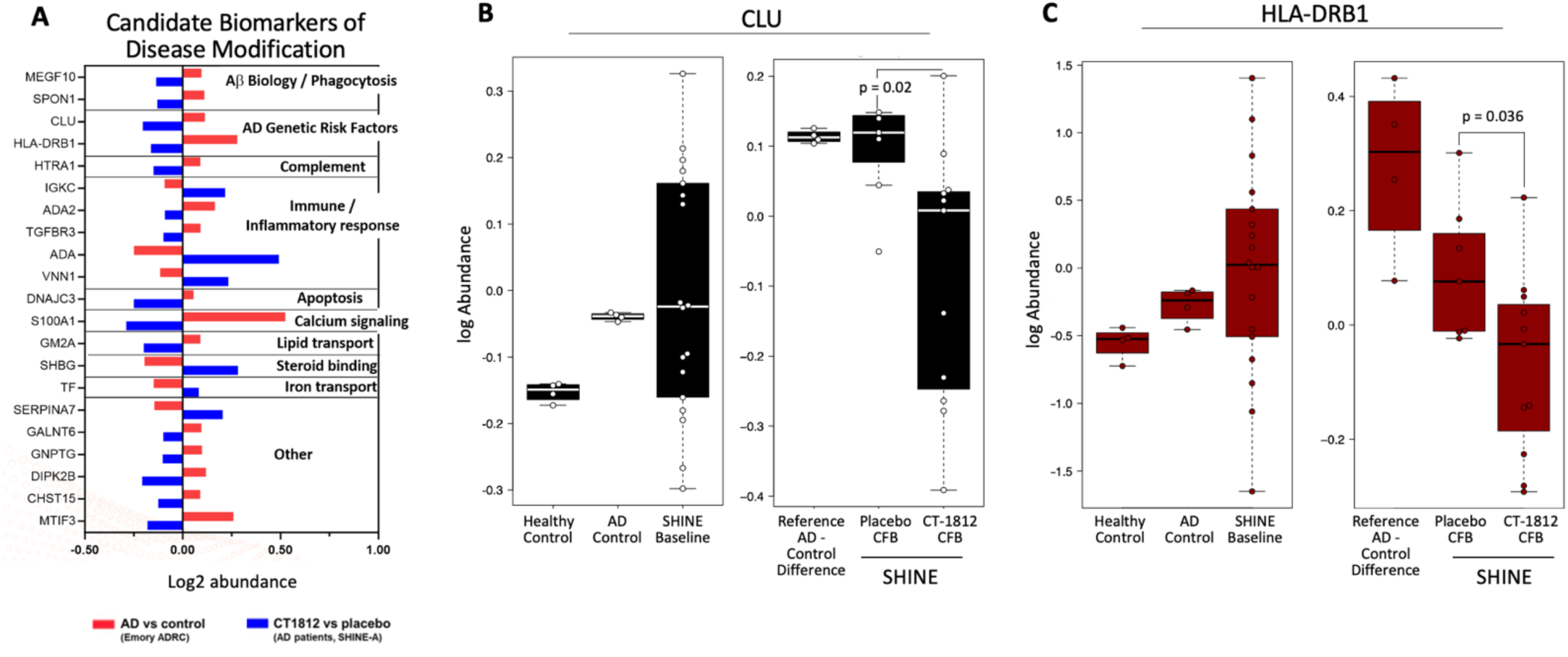
CT1812 pharmacodynamic biomarkers of disease modification. **A.** Forest plot of proteins found to move in the opposition of disease state, by functional category of protein. The log2 change in abundance (p ≤ 0.05) of CSF proteins found to be disrupted in AD (AD vs. healthy control reference standards, p ≤ 0.05, red bars) and significantly normalized (in opposing direction of AD vs. control) by CT1812 (CT1812 vs. placebo, p ≤ 0.05, blue bars) were plotted. CFB = change from baseline. **B.** Clusterin had higher abundance in the pooled AD cohort than in healthy control CSF (left graph). CT1812-treated patient CSF had significantly lower (p = 0.02) clusterin change from baseline abundance than did placebo-treated patient CSF, movement in the opposite direction of disease state. **C**. Likewise, HLA-DRB1 (HLA class II histocompatibility antigen, DRB1 beta chain) abundance was higher in the pooled AD cohort than healthy control CSF (B, left graph), but CT1812-treated patient CSF had significantly lower (p = 0.036) change from baseline abundance than did placebo-treated patient CSF, movement in the opposite direction of disease state. **B-C.** Data is presented as mean (line within box) log abundance +/-SEM (box height) and +/-SD (error bars).

### Impact of CT1812 treatment on Abeta 40 and 42 and correlated proteins

Given the relevance of Aβ to the mechanism of action of CT1812 and to AD, CSF Aβ42 and Aβ40 levels were measured longitudinally in SHINE-A participants using clinically validated assays (Lumipulse; Gobom et al., 2022) (Figure 6A). As this is an exploratory, discovery phase biomarker study to identify possible changes in the CSF proteome following CT1812 treatment, only those samples from participants with confirmed CSF CT1812 levels at the end of study were analyzed. When Aβ42 and Aβ40 change from baseline levels were analyzed in this fashion, both were significantly reduced compared to the change from baseline placebo levels (Aβ42: -104.0 pg/ml for CT1812 vs. 83.4 for placebo, p = 0.0060; Aβ40: - 1594.9 pg/ml for CT1812 vs. 1280.7 pg/ml for placebo, p = 0.0073) (Figure 6A).

**Figure 6.**
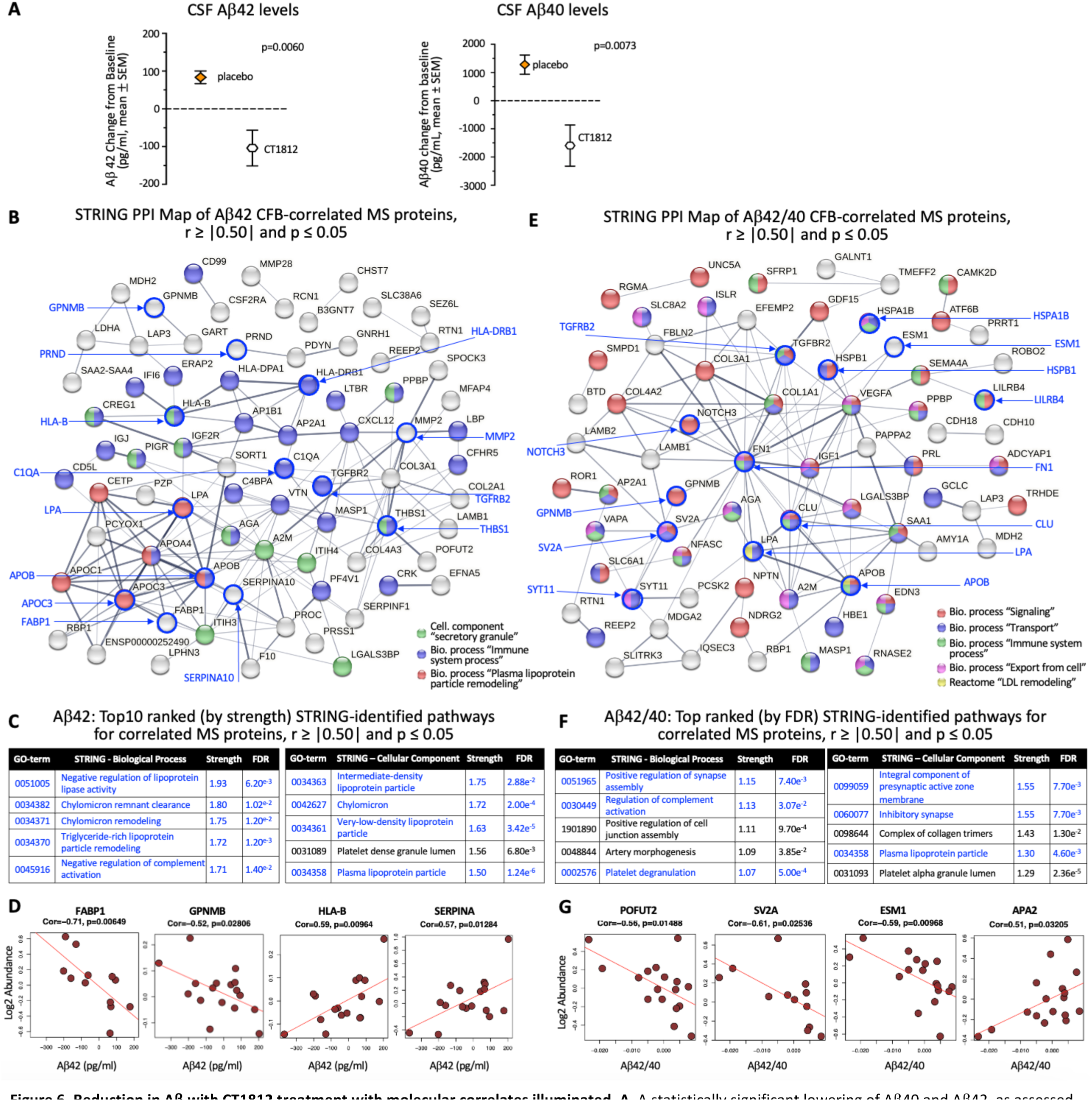
Reduction in Aβ with CT1812 treatment with molecular correlates illuminated. **A.** A statistically significant lowering of Aβ40 and Aβ42, as assessed via Lumipulse assay, was found in CT1812-vs. placebo-treated treatment-compliant patient CSF (n = 7, placebo; n = 12 CT1812). **B**. STRING (v11.5) Protein-Protein Interaction (PPI) map of MS protein CFB levels found to be significantly correlated (r ≥ |0.50| and p ≤ 0.05) with Aβ42 CFB levels (121 proteins; medium confidence, disconnected nodes not shown). Proteins of interest are indicated with royal blue arrows: APOB, Apolipoprotein B; APOC3, apolipoprotein C3; C1QA, Complement C1q subcomponent subunit A; FABP1, fatty acid binding protein 1; GPNMB, Glycoprotein non-metastatic B; HLA-B, major histocompatibility complex, class I, B; HLA-DRB1, HLA class II histocompatibility antigen, DRB1 beta chain; LPA, Lipoprotein(a); MMP2, matrix metallopeptidase 2; PRND, Prion protein 2 (doublet); SERPINA10, serpin family A member 10; TGFRB2, Transforming growth factor, beta receptor II; THBS1, Thrombospondin 1. **C**. Top-ranked (by Strength) Biological Process and Cellular Component terms revealed high representation of lipoprotein-related networks and immune-related pathways (See B and C). **D**. Scatter plots show correlations between Aβ42 CFB and TMT-MS CFB levels for select proteins. Each dot is value from individual participant. **E**. STRING (v11.5) PPI map of MS protein CFB levels found to be significantly correlated (r ≥ |0.50| and p ≤ 0.05) with Aβ42/40 ratio CFB (104 proteins, medium confidence, disconnected nodes not shown). Proteins of interest are indicated with royal blue arrows: APOB; CLU, clusterin; ESM1, endothelial cell specific molecule 1; FN1, fibronectin 1; GPNMB; HSPA1B, heat shock protein 1B; HSPB1, heat shock protein family B (small) member 1; LILRB4, leukocyte immunoglobulin like receptor B4; LPA; NOTCH3, notch receptor 3; SV2A, synaptic vesicle glycoprotein 2a; SYT11, synaptotagmin 11; TGFRB2. **F**. Top-ranked (by Strength) Biological Process and Cellular Component terms revealed high representation of synapse, lipoprotein, and secretion-related networks (also see E). **G**. Scatter plots show correlations between Aβ42/40 CFB and TMT-MS CFB levels for select proteins. APA2, Aspartic proteinase A2; POFUT2, protein O-fucosyltransferase 2. Each dot is value from individual participant.

To gain mechanistic insights and identify proteins associated with the decrease in Aβ42 and Aβ40 levels, Pearson correlation analysis was performed across all proteins detected via TMT-MS against the quantitative assay values. The change from baseline TMT-MS values most strongly correlated with change from baseline Aβ42 values (r ≥ |0.50| and p ≤ 0.05) were identified and included proteins mechanistically linked to Aβ42: apolipoproteins (Kim et al., 2009), complement (Maier et al., 2008) and other inflammatory proteins (Lee et al., 2008) (Figure 6B-C). The protein-protein interaction network was visualized using STRING (Figure 6B). Interestingly, proteins that correlate to the difference or change in Aβ42 levels from baseline showed a very high interconnectivity, with significant GO terms for Cellular Component that included “Secretory granule” (green nodes), and for Biological processes “Immune system process” (blue nodes) and “Plasma lipoprotein particle remodeling” (red nodes). Proteins of interest, highlighted on the protein-protein interaction map in blue circles, include FABP1 and PRND, which were also differentially expressed in participants treated with CT1812 compared to placebo (Figure 4A). Other proteins that correlated with the change in Aβ42 levels that were also differentially abundant following CT1812 treatment (Figure 4A), include JAG2 and DISP3, yet were not on the protein-protein interaction map (data not shown). The top-ranked Biological Process and Cellular Component GO terms for proteins correlated to Aβ levels included lipoprotein-related processes (Figure 6C). In alignment, the top-ranked Reactome term was “LDL remodeling” (data not shown). Scatter plots illustrating the significant correlation between the change from baseline in TMT-MS-detected protein levels and change from baseline Aβ42 for individual participants are in Figure 6D for representative proteins.

We also examined the proteins in the CSF proteome that correlated to the change in Aβ42/40 ratio change from baseline. Although the mean Aβ42/40 ratio change from baseline did not differ between CT1812 and placebo groups (-0.003659 ± 0.002533 SEM for CT1812, 0.001715 ± 0.003571 SEM for placebo, p=0.22), we performed correlation analyses with the change from baseline ratio calculated across participants to all proteins detected via MS-TMT (change from baseline). Similar to the change in strictly Aβ42, the proteins correlated to the change in ratio (r ≥ |0.50| and p ≤ 0.05) were highly interconnected, with notable protein-protein interaction network hub proteins including the genetic AD risk factor CLU, which is decreased in participants receiving CT1812 compared to placebo controls, and the synaptic proteins Syt11 and SV2A (Figure 6E). Other proteins of note, POFUT2 and KRT80 (unconnected nodes hidden from protein-protein interaction map) were correlated with both Aβ42 and Aβ42/40 (data not shown), as well as differentially abundant in CT1812 vs. placebo (Figure 4A). The correlates represented a number of terms relevant to AD and the CT1812 mechanism of action, including the Biological processes “Signaling” (red nodes), “Transport” (blue nodes), “Immune system process” (green nodes), and “Export from cell” (pink nodes), and the Reactome term “LDL remodeling” (yellow nodes). The top ranked (by strength) STRING-identified Biological Process GO and Cellular Component GO terms for Aβ42/40 ratio change from baseline correlates represent synapse assembly, immune function, and secretory processes (Figure 6F). When the Biological Process and Cellular Component GO terms were sorted instead by FDR, secretion- and exocytosis-related elements were strongly represented (Biological Process “Regulated exocytosis” rank 2, “Secretion by cell,” “Export from cell,” and “Secretion” rank 7-9; Cellular Component “Vesicle” rank 3, “Extracellular vesicle,” “Secretory vesicle,” “Extracellular exosome,” and “Secretory granule” rank 5-8; data not shown). Scatter plots illustrating a significant correlation between the change from baseline in MS-detected protein levels and change from baseline Aβ42/40 for individual participants are shown for representative proteins (Figure 6G).

### CT1812 impacted CSF proteins associated with changes in cognitive function

The primary exploratory efficacy measure in SHINE is change from baseline cognitive performance measured by ADAS-Cog11. Although significance in this measure was not expected in the interim analysis of only 24 participants, a clinically meaningful, albeit non-significant, favorable 3-point difference was observed in intended-to-treat participants treated with CT1812 compared to placebo (Figure 7A). To ascertain whether changes in the CSF proteome may be related to the cognitive performance of participants in the clinical trial, the change from baseline levels of proteins measured via TMT-MS were correlated with the change from baseline in ADAS-Cog11 scores. In participants treated with CT1812, a total of 608 TMT-MS-detected CSF proteins were found to be significantly (p ≤ 0.05) correlated with change from baseline ADAS-Cog11 scores (Figure 7B, full blue circle). Of these, 183 were also found to be significantly (p ≤ 0.05) correlated with ADAS-Cog11 in the all-treatments group (i.e., placebo- and CT1812-treated participants, green intersection of two circles). The 183 correlates detected in both CT1812- and placebo-treated CSF samples were removed, and the 425 remaining correlates (i.e., proteins determined to correlate (r ≥ |0.50|) with ADAS-Cog11 only in CT1812-treated participants, blue section of circle, which may represent biomarkers of efficacy), were analyzed using STRING. These biomarkers include APP, APOE, NRXN1, and NRXN2. In the top-ranked (by strength) Biological Process GO terms (Figure 7C, top panel), immune function, synapse assembly, and tau biology are strongly represented. In the top-ranked (by strength) Disease-gene associations (Figure 7C, bottom panel), neurodegenerative disorders, particularly those involving loss of cognition such as AD and Dementia with Lewy Bodies (DLB), are strongly associated, as well as macular diseases. Scatter plots of the AD-related representative protein LTBP1, from the group of 425 correlates analyzed, show that correlation with ADAS-Cog11 is largely driven by drug-treated samples (Figure 7D, right).

**Figure 7.**
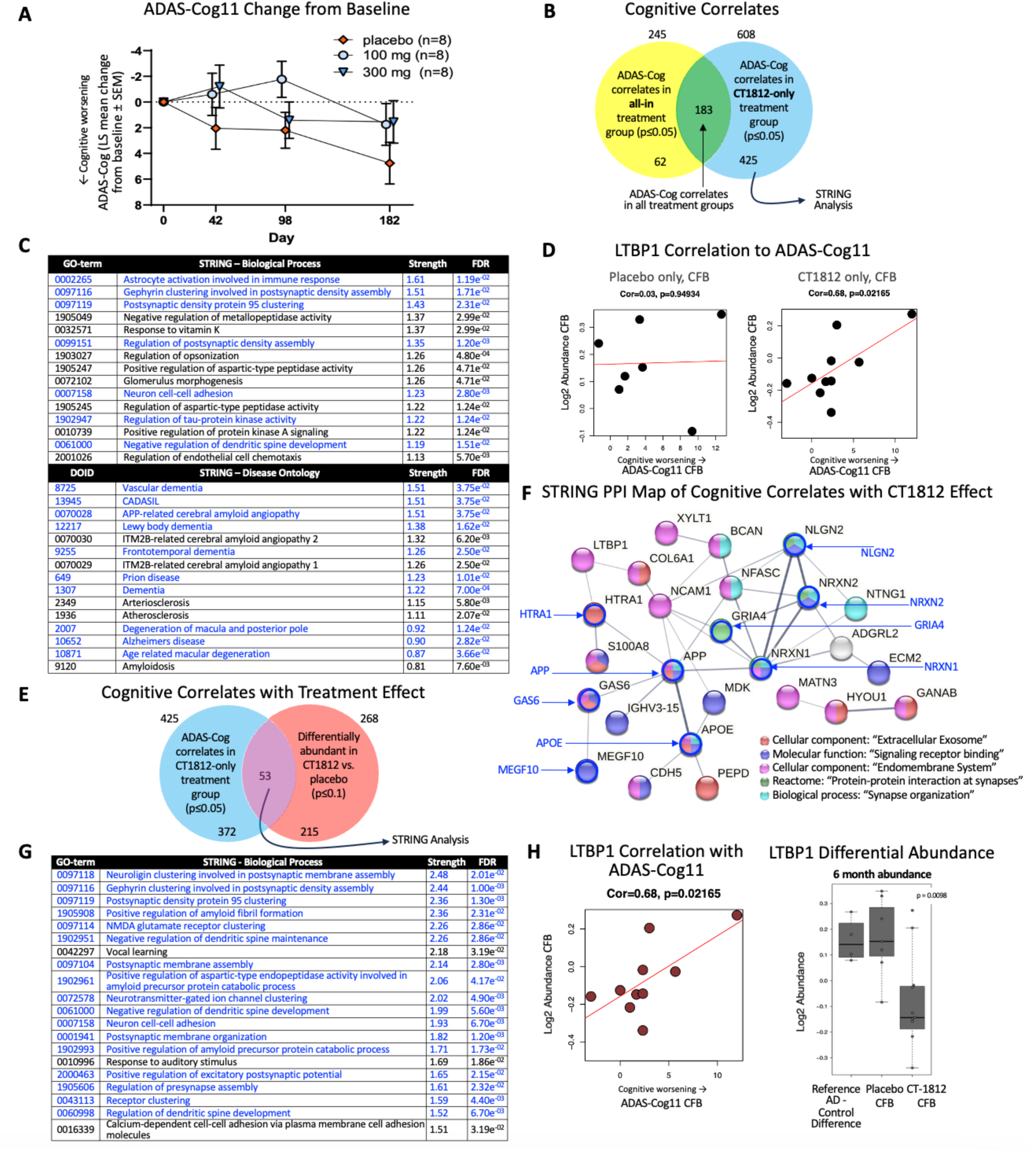
Analysis and identification of molecular correlates with cognitive outcomes. **A.** Change from baseline (CFB) ADAS-Cog11 scores by treatment group over course of 6 months are plotted with all 24 patients (intent to treat). No statistically significant change, as assessed via ANCOVA, due to treatment with CT1812 was observed; a 3 point, clinically meaningful difference was found **B.** To identify molecular correlates with CFB in cognitive function (ADAS-Cog 11) at 6mo, Pearson correlation analysis was performed with CFB levels of proteins assessed via proteomics analysis from a) all patients (i.e., both placebo and CT1812 treated patients; yellow circle) and b) only CT1812-treated patients (blue circle). For the former analysis (a), 245 CSF biomarkers were significantly correlated with ADAS-Cog11 CFB (yellow circle). In the CT1812-treated group only (b), a total of 608 proteins were found to be correlated with ADAS-Cog11 CFB (blue circle). Of these, 183 were also found correlated in the all-treatment group (green overlap). The 183 correlates detected in both CT1812- and placebo-treated CSF samples were removed, and the 425 remaining CT1812 group correlates (i.e., proteins determined to correlate with ADAS-Cog11 only in CT1812-treated patients, outer portion of blue circle) were analyzed using STRING (v12.0). Top-ranked (by Strength) Biological Process GO terms (top panel) and Disease-Gene Associations (bottom panel) are shown (**C**). **D.** Scatter plots showing individual patient LTBP1 log2 abundance CFB and ADAS-Cog11 CSF values are shown for Placebo treated patients only (left) and CT1812-treated only (right), illustrating the correlation is driven by CT1812 and thus this may be a cognitive correlate attributable to a CT1812-specific mechanism of action. **E.** Of the 425 proteins identified that are correlated with ADAS-Cog in the CT1812 group only (blue circle), 53 (purple overlap) were determined to also be differentially abundant in CT1812 vs. placebo CSF (Figure 4A). **F.** These 53 proteins were analyzed using STRING (v12.0) and the Protein-Protein Interaction (PPI) map is shown. Proteins of interest are indicated with royal blue arrows: NLGN2, neuroligin 2; NRXN1 and 2, neurexin 1 and 2; GRIA4, glutamate ionotropic receptor AMPA type subunit 4; APOE, apolipoprotein E; APP, amyloid beta precursor protein; MEGF10, multiple EGF like domains 10; GAS6, growth arrest specific 6; HTRA1, HtrA serine peptidase 1. **G.** Top-ranked (by Strength) Biological Process GO terms. **H.** Correlation of LTBP1 with AGAS-Cog11 in individual CT1812-treated patients (left; each dot is CFB value from individual participant; differential log2 abundance of LTBP1 in AD vs. control CSF, and in SHINE placebo- or CT1812-treated CSF at 6 months. CFB = change from baseline.

To query which of the ADAS-Cog11-correlated proteins were also differentially abundant in participants treated with CT1812 vs. placebo, and therefore presumably impacted by drug treatment, these 425 proteins (Figure 7E, full blue circle), were compared to the proteins found to be significantly (p ≤ 0.10) increased or decreased with drug treatment (Figure 7E, full red circle, 268 proteins), and 53 overlapping correlates were identified (Figure 7E, overlap of blue and red circles). These 53 candidate proteins that correlated with both change in cognition and CT1812 treatment may therefore reflect disease modification and mechanism(s) of action of target engagement. To this end, these proteins were analyzed using STRING, and the resulting protein-protein interaction map shows a number of relevant centralized nodes including APP, APOE, Neurexin 1 and 2 (NRXN1 and NRXN2), and Neuroligin 2 (Figure 7F, red arrows). Synaptic and exosome/endosome components, consistent with the CT1812 mechanism of action, are strongly represented on the protein-protein interaction map. Likewise, the top-ranked (by Strength) Biological Process GO terms largely comprise synapse-related terms (Figure 7G). Scatter plot of the AD-related protein LTBP1, representative from the group of 53 correlates analyzed, shows its negative correlation with ADAS-Cog11 in individual samples (Figure 7H). The box plot shows the differential abundance of LTBP1 in AD control CSF, placebo-treated SHINE participants CSF, and CT1812-treated CSF, demonstrating that LTBP1 was reduced with drug treatment.

## DISCUSSION

In this exploratory interim analysis of the SHINE mild-moderate AD cohort, quantitative proteomics identified pharmacodynamic biomarkers of CT1812 that were pathway engagement biomarkers (i.e., tied to CT1812’s mechanism of action) and disease modification biomarkers (i.e., associated with favorable changes in cognition by CT1812 or normalized towards control levels by CT1812). Brain network mapping, Gene Ontology, and pathway analyses revealed an impact of CT1812 on disease-relevant pathways and biology including synaptic function, lipoprotein and Aβ biology, and immune function. Further, the longitudinal data was leveraged to shed light on proteins and pathways linked to hallmarks of disease such as Aβ, NfL, tau, and Syt1.

### Biomarker discovery

The measurement of CSF biomarkers in clinical trials is imperative to clinical development success, in the selection and recruitment of the appropriate patients, in dose selection, and in the monitoring of changes during treatment. Examining pharmacodynamic biomarkers longitudinally in trials can shed light on whether and how a therapeutic is impacting the processes of disease, confirm the therapeutic mechanism of action, enable the identification of biomarkers that may denote responders or patients who are most likely to benefit from treatment, and reveal surrogate biomarkers of disease modification.

The field of AD biomarker discovery has been challenged by a lack of quantitative, sensitive, specific, and reliable methodologies beyond a small, but growing, set of AD core biomarkers. While examination of the core AD biomarkers including Aβ and tau species is important because they are hallmarks of AD, there are thousands of proteins that are expressed in the brain and also found in CSF. In this study, we have utilized a recently improved quantitative method of MS, TMT paired with benchmark reference standards, facilitating reliable detection of >2,000 proteins including proteins of low abundance, that older methods lacked the sensitivity to detect (Ping et al., 2018), in our clinical trial samples. In this manner, CSF proteomics can provide a comprehensive, quantitative, holistic view of the proteins and pathways impacted in brain in AD and with therapeutic intervention.

In this study, we validated the precision of the quantitation of this method through comparison of the absolute levels of TMT-MS-measured proteins within each sample to those measured using independent, highly quantitative methods that had been clinically validated, finding robust and highly significant correlations for several of the core AD biomarkers including t-Tau, Nrgn, Syt1, and NfL. Assessments of these highly correlated proteins to determine their related biological processes further validated this method and provided rich data from which biological insights can be gleaned. Notably, proteins correlated with the axonal cytoskeletal protein NfL were found to be involved in processes already known to be associated to NfL biology, including axonogenesis and neuron projection development. Taken together with similar findings for qTau, qNrgn, and qSyt1, these results validate the use of TMT-MS proteomics as a highly quantitative method. Furthermore, the novel proteins and pathways we found to be associated with the seminal core AD biomarkers can be leveraged by the field to explore the molecular dynamics of AD.

TMT-MS analysis of reference standards using pooled CSF samples from known AD biomarker positive and negative cohorts revealed more than 500 differentially expressed proteins that were significantly (FDR p ≤ 0.05) differentially abundant (increased or decreased) in AD vs. healthy CSF, and corroborate findings from previous studies (e.g., upregulation of tau, CLU, NfL and NfM; downregulation of Neurexin and APOE) (Bergström et al., 2021; Bertrand et al., 1995; Blennow et al., 1994; Higginbotham et al., 2020; Hölttä et al., 2015). Mapping of these proteins to previously-established functional modules (Johnson et al., 2022) revealed, at the proteome-wide level, that the proteins most impacted by AD are involved in immune processes, metabolism, and neuronal synapses - biological processes and functions known to be disrupted in AD - further supporting the quantitative use of TMT-MS.

In a novel application of this proteomics method, we utilized the AD and control CSF reference standard proteomes as benchmarks for the characterization of the SHINE-A participant population baseline proteome. For key AD-related proteins from the most highly impacted modules, SHINE-A participant CSF levels closely resemble levels in the AD reference cohort and appeared dissimilar to those of the control reference cohort, indicating that the SHINE-A cohort studied in the interim analysis is prototypical of AD. For example, SHINE-A participants had similar levels of elevated MAPT (tau), CLU, NEFL, and NEFM, and similarly reduced levels of APOE and neurexin 3. The relevance of the SHINE population at baseline to patients with AD overall encouragingly supports the possibility that results from this clinical trial may later be replicated in subsequent trials in mild-moderate AD patients, and informs the design of future trials.

### Pharmacodynamic biomarkers of CT1812

Three main categories of candidate pharmacodynamic biomarkers for CT1812 were identified in this study: those reflecting pathway engagement, those reflecting disease modification, and those reflecting both (Figure 8). Comparison of the CT1812- and placebo-treated CSF proteomes revealed pharmacodynamic biomarkers that were significantly altered with drug treatment. These differentially expressed proteins belonged primarily to the Brain Map modules “Synapse/neuron,” “Oligo/myelination,” “MHC complex/immune,” and “Matrisome,” indicating an impact on pathways relevant to AD pathophysiology. In support, many of these brain modules were among the top brain modules dysregulated in AD vs. control (refer to Figure 3), suggesting an impact of CT1812 on disease-related biological processes.

**Figure 8.**
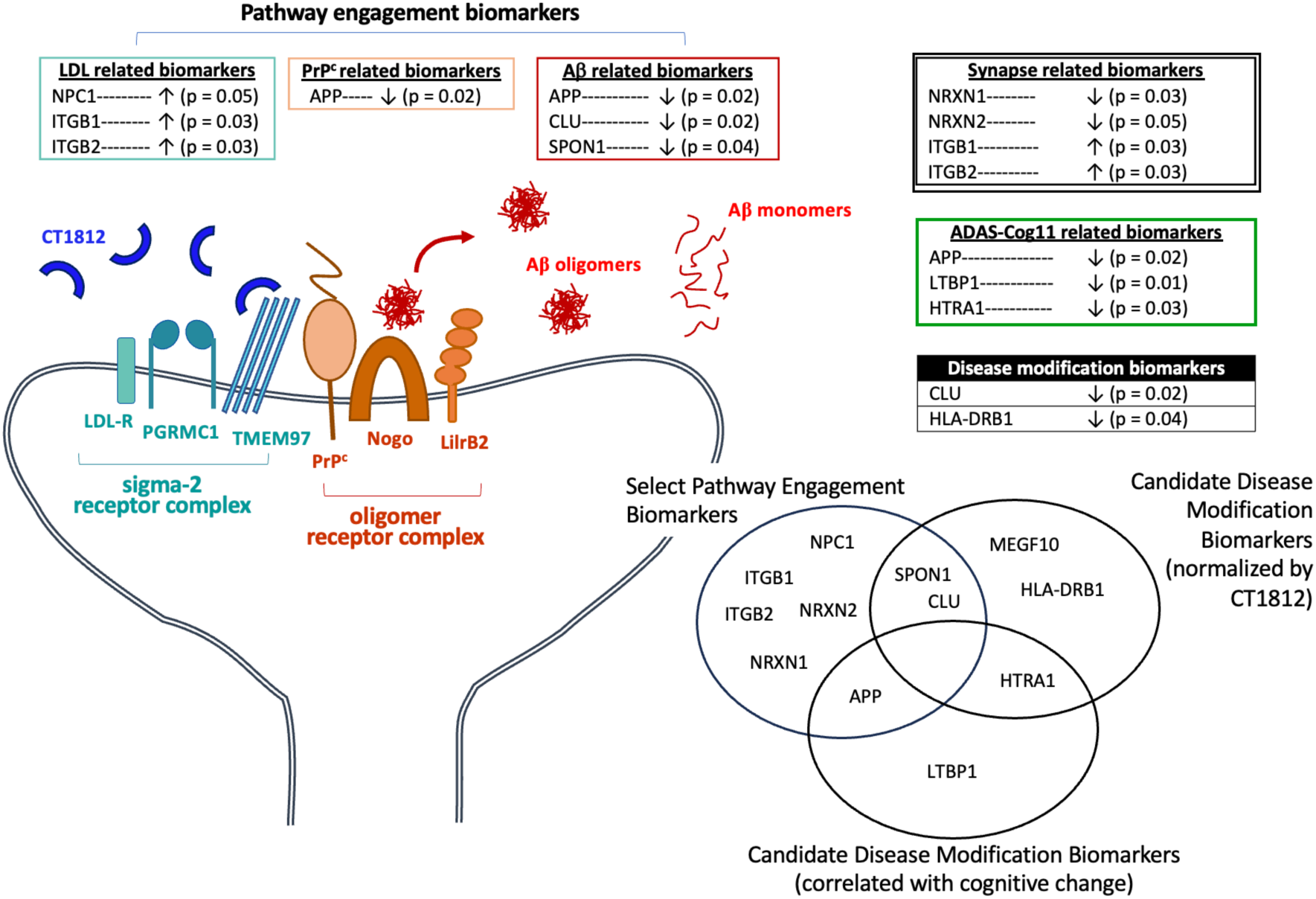
Candidate PD Biomarkers of CT1812: Support for the hypothesized CT1812 mechanism of action. CT1812 (royal blue) binds the sigma-2 receptor (S2R), also known as TMEM97. TMEM97 is localized on the membrane in a complex (turquoise) with the LDL receptor (LDLR) and PGRMC1. Aβ oligomers (red) bind the oligomer receptor complex (orange), which comprises PrP^c^, Nogo, and LilrB2. Binding of CT1812 to the S2R complex induces a conformational change in the oligomer receptor complex, disengaging the oligomer from the synapse (Izzo et al., 2021). Clockwise from bottom left: several biomarkers identified in the SHINE interim CSF proteomics analysis suggest CT1812 impacts synapses (NRXN1, NRNX2, ITGB1, ITGB2; double lined box). NPC1, ITGB2, and ITGB1 (turquoise box) suggest the LDLR may be impacted by CT1812 treatment- and these biomarkers may reflect target engagement. In addition, APP reduction is consistent with a PrP^c^-related mechanism of action (orange box). Reductions in APP, CLU, and SPON1 (red box) are consistent with a negative regulation of Aβ, and may also reflect pathway engagement. CLU and HLA-DRB1 (black box), which were increased in abundance in AD vs. control CSF but decreased in abundance in CT1812 vs. placebo-treated patient CSF, are two candidate biomarkers of disease-modification. APP, LTBP1, and HTRA1 were identified as molecular correlates of ADAS-Cog and may also be candidate surrogate biomarkers of efficacy.

A number of the individual proteins impacted by CT1812 are known to be genetically linked to AD, including APP and CLU, both reduced with CT1812 treatment. Specific mutations in APP, the precursor protein for all forms of Aβ (monomer, oligomer, proto-fibril, fibril and plaque forms), lead to autosomal dominant familial AD while other mutations in APP confer an increased or decreased risk of AD in cases of non-familial, sporadic AD (Cruchaga et al., 2012; Kunkle et al., 2019). CLU, known to be elevated in AD (Shepherd et al., 2020), is a genetic risk factor with a single nucleotide polymorphism (SNP) conferring increased risk for sporadic, or late-onset AD (LOAD) (Harold et al., 2009; Lambert et al., 2009). Another protein found to be regulated by CT1812 was SPON1, which binds to APP and inhibits its cleavage by β-secretase, and has been associated with risk of dementia and AD (Jahanshad et al., 2013; Sherva et al., 2014). Of note, the proteins contributing to “Positive regulation of amyloid fibril formation” were APP, CLU, and SPON1, all of which were decreased in abundance in CT1812 vs. placebo (Figure 4A). The ability of CT1812 to impact levels of these proteins is therefore suggestive that CT1812 is modifying disease-relevant pathways. Given both the disease relevance and the mechanism of action of CT1812 in impacting Aβ dynamics in displacing Aβ oligomers from neuronal synapses, it is encouraging that a large number of the identified proteins (including APP, CLU, and APL2) relate to Aβ.

Many of the identified pharmacodynamic biomarkers can also be considered candidate biomarkers of CT1812 pathway engagement, pending confirmation or replication in independent cohorts of AD patients treated with CT1812. CLU is an apolipoprotein (APOJ) that binds Aβ with high affinity (Beeg et al., 2016) and plays a role in sequestering Aβ oligomers, prevents the oligomerization of Aβ, and impacts Aβ clearance through a number of potential mechanisms, including astrocyte uptake and clearance into the ISF (Foster et al., 2019; Narayan et al., 2011). Because Aβ oligomer displacement and clearance into the interstitial fluid (ISF) and CSF is a central mechanism of CT1812 (Izzo et al., 2021; LaBarbera et al., 2023), CLU is a potential biomarker of CT1812 pathway engagement. CLU is also an astrocyte-derived protein and has been shown to impact Aβ oligomer uptake by astrocytes (Nielsen et al., 2010); astrocyte activation has been recently linked to CT1812 (Colom-Cadena M et al., 2023). Further mechanistic studies of both CT1812 and CLU will be needed to determine whether the precise mechanisms of Aβ oligomer engagement are related.

Another finding that provides proof of mechanism for CT1812 is the lower change from baseline levels of the amyloid precursor protein (APP) in CT1812 vs. placebo. CT1812 acts through S2R, also known as TMEM97, which itself directly interacts with prion protein (PRNP; also known as PrP^c^). APP is known to be regulated by changes in cellular PrP^c^ mediated activity and localization to lipid rafts (Parkin et al., 2007), and silencing of PrP^c^ has been shown to reduce APP processing by β-secretase and decrease levels of Aβ40 and Aβ42 through a (Pietri et al., 2013). Thus, it is possible that when CT1812 binds the S2R, resulting in the displacement of Aβ oligomers from the oligomer receptor, this leads to the observed downstream change in APP levels due to altered PrP^c^ -mediated signaling. Reduced APP levels may therefore be an indication that the S2R/ PrP^c^ receptor complex has been functionally engaged.

S2R is also thought to complex with the low-density lipoprotein receptor (LDLR) (Riad et al., 2018, 2020). NPC1 (Niemann-Pick disease type C1), ITGB1 (β1 integrin), and ITGB2 (β2 integrin), all of which are associated with the GO term “Cellular response to low-density lipoprotein particle stimulus” and impact LDL cholesterol intracellular trafficking (Hoque et al., 2015), had higher change from baseline levels with CT1812 treatment compared to placebo. Furthermore, by detecting extracellular changes and responsively regulating synapse formation through actin remodelling, integrins play a role in neuronal synaptic assembly and plasticity (Y. K. Park & Goda, 2016), also relevant to the CT1812 mechanism of action. If these pharmacodynamic biomarker findings are replicated by larger clinical trials, these candidate pathway engagement biomarkers could be used to determine whether individual patients are responding to treatment, or have sufficient drug exposure, and may help to further refine dose selection in future trials to foster the clinical development success of CT1812.

Finally, findings from this study also support the continued development of CT1812 for additional indications. Disease-gene associations identified by STRING for proteins found to correlate with changes in cognitive function included DLB. A phase 2 clinical trial to study the safety and efficacy of CT1812 in subjects with mild to moderate DLB (NCT05225415) is currently recruiting. In addition, one biological process found to be impacted by differentially abundant proteins by Metacore pathway analysis was “Aberrant lipid trafficking in metabolism in age-related macular degeneration (AMD)” (Figure 4E), and two disease ontology associations identified by STRING were “Age related macular degeneration” and “Degeneration of macula and posterior pole” (Figure 7C). An ongoing clinical trial is currently testing the hypothesis that CT1812 will impact dry AMD (NCT05893537). In this way, biomarker findings gleaned from clinical trials in AD may help inform the design and analysis of studies of other related indications.

### New learnings to shed light on mechanisms related to CSF amyloid beta levels

CSF Aβ42 and Aβ40 levels, including calculation of the Aβ42/40 ratio, are routinely assessed to evaluate impact on disease biology and for diagnosis of AD (Ashton et al., 2022). In the present study, analysis of change from baseline levels of Aβ42 and Aβ40 in participants who were actively taking their treatment (as indicated via bioanalysis of drug exposure levels) using the clinically validated Lumipulse assays, were significantly lower in CT1812 compared to placebo treated participants. This finding is consistent with the known impact of CT1812 on Aβ biology from preclinical (Izzo et al., 2021) and clinical studies (LaBarbera et al., 2023) (impact on Aβ through PrP^c^, as described above), albeit the first evidence of a potential impact on monomeric Aβ levels in CSF. Given CT1812’s mechanism of action and that Aβ is the chief hallmark of AD, a clinically validated target for AD for which there are disease modifying, Aβ-lowering therapies available to patients (Budd Haeberlein et al., 2022; van Dyck et al., 2023), we leveraged our TMT-MS-detected protein dataset to ask whether other meaningful proteins shifted in concordance with Aβ42 or Aβ42/40, as well as with CT1812 treatment. In CT1812-treated CSF, Aβ42 change from baseline levels were significantly correlated with change from baseline abundances of 121 proteins. GO term and Reactome analysis of these proteins using STRING revealed a high relevance of these proteins to lipoprotein (LDL, chylomicron, phospholipid, etc.) biology. Likewise, the change from baseline Aβ42/40 ratio was found to be significantly correlated with proteins relevant to synapse assembly. As the LDL receptor is thought to interact in a complex with CT1812 target receptor S2R (Riad et al., 2018, 2020), and as synapses are the mechanistic target of CT1812 (Izzo et al., 2021), these correlations are supportive of the CT1812 mechanism of action and suggest a role for CT1812 in Aβ biology. Additionally, although Aβ40 and Aβ42 peptides were not assessed via TMT-MS, these Lumipulse-detected reductions in Aβ40 and Aβ42 are consistent with the decrease in APP, the precursor protein for Aβ40 and Aβ42 generation, detected via MS in CT1812 vs. placebo (Figure 4A, C). We queried whether the decrease in APP protein levels could be driven by a lowering of the Aβ40-42 peptide and examined the peptide level proteomics data. However, the decrease in APP spanned the entire protein and not just the peptides comprising the 1-40/1-42 amino acid sequence (data not shown). Future investigation of this finding is warranted to assess replication of findings in an independent clinical cohort and to better understand the precise molecular mechanisms underlying this effect.

### Identification of candidate surrogate biomarkers of disease modification

Our study also identified biomarkers of disease modification: proteins that are aberrant in AD but that were normalized with CT1812 treatment (i.e., significantly changed in the opposite direction of disease progression). CLU was elevated in AD and in SHINE-A baseline samples compared to healthy control but reduced in CT1812 vs. placebo CSF after 6 months of treatment, indicative of CLU as both a candidate biomarker of CT1812 pathway engagement and a candidate pharmacodynamic biomarker. HLA-DRB1, a histocompatibility gene that has been identified by GWAS as a risk factor for AD (Lu et al., 2017), was also found to shift towards healthy control levels with CT1812 treatment. Other proteins that likewise moved towards normalization are known to relate to AD (e.g., MEGF10, SPON1), or to meaningful biological processes (e.g., immune/ inflammatory responses and lipid transport). The impacted proteins support proof of mechanism that CT1812 impacts synapse, inflammation, secretion, and amyloid biology. These findings not only support the disease-relevance of changes induced by CT1812 but are also preliminary evidence for potential surrogate biomarkers to use in future clinical trials and for treatment purposes.

Finally, the goal for CT1812 and therapeutics aimed at treating AD is, ultimately, a clinically meaningful impact on cognitive status. Cognitive performance, as measured by ADAS-Cog11, is the primary exploratory efficacy readout in the SHINE trial. This interim analysis of the first 24 participants assessed in the SHINE trial showed a clinically meaningful, albeit statistically non-significant, 3-point improvement in ADAS-Cog11. As with Aβ above, we leveraged the TMT-MS proteome dataset collected from these participants to ask whether other meaningful proteins shifted in concordance with cognitive performance. Correlation analyses of MS-measured proteins change from baseline in CT1812-treated CSF and cognitive outcomes, as measured using the ADAS-Cog11 score change from baseline, revealed 425 proteins that were significantly correlated (p ≤ 0.05). The correlates that had also shifted in the all-treatment cohort, which includes placebo-treated participants, were removed to ensure the analysis did not consider proteins that shifted over the course of the trial even without treatment. The remaining 53 cognitive correlates that had also been differentially expressed in CT1812-vs. placebo-treated CSF are pharmacodynamic biomarkers of CT1812 that may also represent disease modification. Gene ontology analysis of these proteins using STRING revealed a number of terms related to the CT1812 mechanism of action, including astrocyte activation (Colom-Cadena M et al., 2023) and synapse assembly. STRING-identified diseases associated with the cognition-correlated proteins were largely dementia-related. 14 of the top 20 ranked GO Biological Processes impacted were related to synapses, with another three relating to amyloid biology. Together, these findings corroborate the preclinical evidence that CT1812 is protective of synapses (Izzo et al., 2021), and suggest the proteomic shifts associated with CT1812 treatment may relate to cognitive status.

## Conclusion

The proteomics analyses reported in this study suggest that the S2R modulator CT1812 may be a promising disease-modifying treatment for AD. This unbiased approach to identifying biomarkers associated with CT1812 treatment revealed changes to synapse, immune, secretion, and lipoprotein-related biological processes, supporting the purported mechanism of action of CT1812 and indicating relevance to AD pathophysiology (Figure 8). Identifying these changes through unbiased methods underscores the clinical translatability of mechanisms investigated in previous preclinical hypothesis-based research of CT1812 and S2R modulating compounds (Izzo et al., 2014, 2021). Pharmacodynamic biomarkers of CT1812 pathway engagement and disease modification were identified. If these results are replicated by larger studies, these biomarkers have the potential to support clinical trials and patient care in the future. This interim exploratory biomarker analysis enabled the establishment of a framework and analysis pipeline to use for our future biomarker assessments of the full SHINE trial and other clinical trials underway. Analysis of the full SHINE trial of ∼150 participants will be important to assess replication of findings and confirm the hypothesized pharmacodynamic effect of CT1812 in AD patients.

## Acknowledgements

The authors thank Evi Malagise and Lora Waybright for technical assistance in earlier analyses.

This work was supported by funding from the National Institute on Aging (1R01AG058660-01) and the Alzheimer’s Drug Discovery Foundation, and by Cognition Therapeutics.

