## Supplemental Figure 1 for "An interim exploratory biomarker analysis of a Phase 2 clinical trial to assess the impact of CT1812 in Alzheimer’s disease"

BN Lizama[1] et al.

### SUPPLEMENTAL MATERIALS

**A** qNfL:NEFM MS Correlation

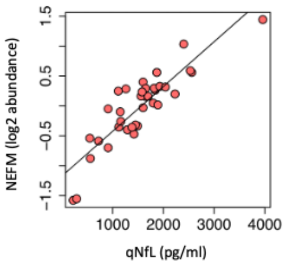


**B** STRING: Top Pathways Identified*

| ELISA Trait | Biological Process GO Term (Top-ranked by FDR) | Molecular Function GO Term (Top-ranked by FDR) |
|---|---|---|
| Tau | 1. Cell adhesion ($5.12e^{-23}$)<br>2. Response to stimulus ($2.23e^{-19}$)<br>3. Nervous system development ($1.25e^{-17}$) | 1. Signaling receptor binding ($3.76e^{-13}$)<br>2. Protein binding ($1.60e^{-12}$)<br>3. Endopeptidase inhibitor activity ($2.45e^{-09}$) |
| NfL | 1. Negative regulation of cell morphology in differentiation ($6.77e^{-05}$)<br>2. Regulation of cell morphology in differentiation ($6.77e^{-05}$)<br>3. Regulation of axonogenesis (0.00011) | 1. Chemorepellent activity (0.0029) |
| Nrgn | 1. Cell adhesion ($1.09e^{-31}$)<br>2. Response to stimulus ($1.30e^{-21}$)<br>3. Nervous system development ($1.27e^{-20}$) | 1. Signaling receptor binding ($4.94e^{-19}$)<br>2. Protein binding ($2.03e^{-16}$)<br>3. Cell adhesion molecule binding ($7.72e^{-13}$) |
| Syt1 | 1. Cell adhesion ($1.95e^{-27}$)<br>2. Nervous system development ($8.19e^{-18}$)<br>3. System development ($8.19e^{-18}$) | 1. Calcium ion binding ($8.12e^{-14}$)<br>2. Protein binding ($1.04e^{-12}$)<br>3. Signaling receptor binding ($2.38e^{-12}$) |

*Ranked by FDR. STRING pathway analysis performed on MS proteins correlated (r) ≥ |0.50| and p ≤ 0.01; Genes named "0" removed for STRING. GO, Gene Ontology; FDR, False Discovery Rate; Top 6 ranked pathways are displayed; Pathways of biological interest are highlighted in royal blue.

**C** STRING PPI Map of qNrgn MS Correlated proteins [(r) ≥ |0.70|, p ≤ 0.01]

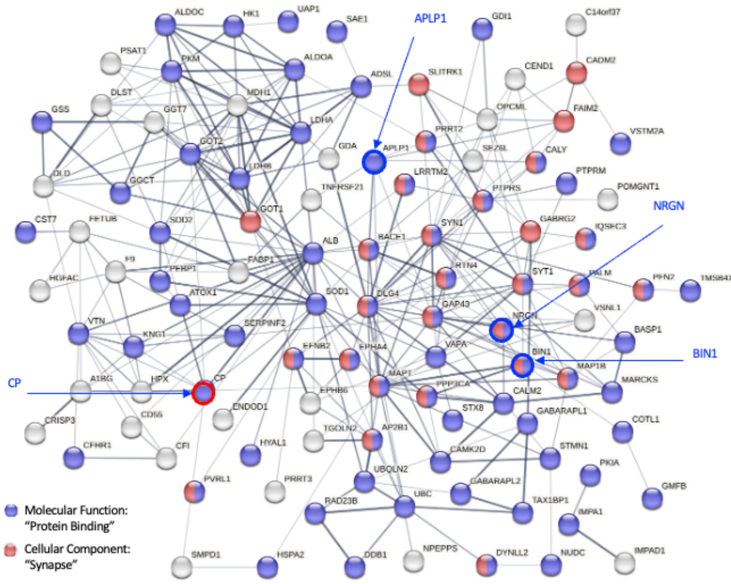


**D** Top ranked Cellular Component pathways for qNrgn MS correlated proteins [(r) ≥ |0.70|, p ≤ 0.01]

| GO Term | STRING - Cellular Component | Strength | FDR^ |
|---|---|---|---|
| 0031982 | Vesicle | 0.45 | $3.29e^{-16}$ |
| 0070062 | Extracellular exosome | 0.51 | $1.28e^{-10}$ |
| 0005576 | Extracellular region | 0.37 | $1.28e^{-10}$ |
| 0098984 | Neuron to neuron synapse | 0.88 | $4.16e^{-09}$ |
| 0005615 | Extracellular space | 0.40 | $4.16e^{-09}$ |
| 0098794 | Postsynapse | 0.73 | $1.72e^{-08}$ |
| 0030424 | Axon | 0.71 | $1.01e^{-07}$ |
| 0014069 | Postsynaptic density | 0.84 | $3.89e^{-07}$ |
| 0031410 | Cytoplasmic vesicle | 0.42 | $3.89e^{-07}$ |
| 0045202 | Synapse | 0.51 | $2.60e^{-06}$ |
| 0099503 | Secretory vesicle | 0.55 | $7.98e^{-06}$ |
| 0072562 | Blood microparticle | 1.07 | $1.44e^{-05}$ |
| 0098978 | Glutamatergic synapse | 0.76 | $1.99e^{-05}$ |
| 0071944 | Cell periphery | 0.25 | $2.46e^{-05}$ |
| 0005886 | Plasma membrane | 0.25 | $2.52e^{-05}$ |
| 0043005 | Neuron projection | 0.47 | $3.17e^{-05}$ |
| 0097060 | Synaptic membrane | 0.73 | $3.65e^{-05}$ |
| 0030054 | Cell junction | 0.39 | $5.10e^{-05}$ |
| 0016020 | Membrane | 0.16 | $9.60e^{-05}$ |
| 0042995 | Cell projection | 0.36 | $1.50e^{-04}$ |
| 0098685 | Schaffer collateral – CA1 synapse | 1.10 | $1.60e^{-04}$ |
| 0120025 | Plasma membrane bounded cell projection | 0.36 | $1.60e^{-04}$ |
| 0060205 | Cytoplasmic vesicle lumen | 0.74 | $1.80e^{-04}$ |
| 0031091 | Platelet alpha granule | 1.06 | $2.30e^{-04}$ |
| 0045211 | Postsynaptic membrane | 0.77 | $2.30e^{-04}$ |
| 0030659 | Cytoplasmic vesicle membrane | 0.54 | $2.60e^{-04}$ |
| 0036477 | Somatodendritic compartment | 0.52 | $2.60e^{-04}$ |
| 0005737 | Cytoplasm | 0.12 | $2.90e^{-04}$ |
| 0098839 | Postsynaptic density membrane | 1.02 | $3.50e^{-04}$ |
| 0030427 | Site of polarized growth | 0.84 | $4.20e^{-04}$ |
| 0031093 | Platelet alpha granule lumen | 1.12 | $4.40e^{-04}$ |
| 0044297 | Cell body | 0.58 | $5.10e^{-04}$ |
| 0034774 | Secretory granule lumen | 0.71 | $6.40e^{-04}$ |
| 0031012 | Extracellular matrix | 0.60 | $6.40e^{-04}$ |
| 0030141 | Secretory granule | 0.50 | $6.70e^{-04}$ |

**E** Top ranked Cellular Component pathways for qSyt1 MS correlated proteins [(r) ≥ |0.70|, p ≤ 0.01]

| GO Term | STRING - Cellular Component | Strength | FDR^ |
|---|---|---|---|
| 0099056 | Integral component of presynaptic membrane | 1.36 | $7.00e^{-03}$ |
| 0005794 | Golgi apparatus | 0.53 | $7.00e^{-03}$ |
| 0070062 | Extracellular exosome | 0.47 | $7.00e^{-03}$ |
| 0031982 | Vesicle | 0.37 | $7.00e^{-03}$ |
| 0012505 | Endomembrane system | 0.31 | $7.00e^{-03}$ |
| 0016020 | Membrane | 0.20 | $7.00e^{-03}$ |
| 0098978 | Glutamatergic synapse | 0.86 | $1.01e^{-02}$ |
| 0098984 | Neuron to neuron synapse | 0.83 | $1.27e^{-02}$ |
| 0005791 | Rough endoplasmic reticulum | 1.23 | $1.47e^{-02}$ |
| 0097060 | Synaptic membrane | 0.81 | $1.47e^{-02}$ |
| 0045202 | Synapse | 0.52 | $1.47e^{-02}$ |
| 0098793 | Presynapse | 0.72 | $1.56e^{-02}$ |
| 0005912 | Adherens junction | 0.99 | $1.83e^{-02}$ |
| 0030054 | Cell junction | 0.42 | $2.40e^{-02}$ |
| 0005788 | Endoplasmic reticulum lumen | 0.82 | $2.81e^{-02}$ |
| 0005576 | Extracellular region | 0.30 | $2.81e^{-02}$ |
| 0032584 | Growth cone membrane | 1.94 | $3.09e^{-02}$ |
| 0014069 | Postsynaptic density | 0.81 | $3.09e^{-02}$ |
| 0098794 | Postsynapse | 0.65 | $3.19e^{-02}$ |

**F** Top ranked Cellular Component pathways for qTau MS correlated proteins [(r) ≥ |0.70|, p ≤ 0.01]

| GO Term | STRING - Cellular Component | Strength | FDR^ |
|---|---|---|---|
| 0031982 | Vesicle | 0.49 | $6.17e^{-12}$ |
| 0070062 | Extracellular exosome | 0.56 | $9.94e^{-08}$ |
| 0005576 | Extracellular region | 0.38 | $4.74e^{-06}$ |
| 0005615 | Extracellular space | 0.41 | $4.08e^{-05}$ |
| 0005829 | Cytosol | 0.31 | $4.75e^{-05}$ |
| 0030424 | Axon | 0.74 | $4.86e^{-05}$ |
| 0098794 | Postsynapse | 0.74 | $4.86e^{-05}$ |
| 0030659 | Cytoplasmic vesicle membrane | 0.69 | $5.51e^{-05}$ |
| 0031410 | Cytoplasmic vesicle | 0.44 | $1.10e^{-04}$ |
| 0005737 | Cytoplasm | 0.16 | $1.10e^{-04}$ |
| 0043005 | Neuron projection | 0.55 | $1.30e^{-04}$ |
| 0045202 | Synapse | 0.53 | $4.50e^{-04}$ |
| 0098805 | Whole membrane | 0.47 | $8.00e^{-04}$ |
| 0120025 | Plasma membrane bounded cell projection | 0.43 | $8.00e^{-04}$ |
| 0098978 | Glutamatergic synapse | 0.80 | $1.30e^{-03}$ |
| 0098984 | Neuron to neuron synapse | 0.79 | $1.50e^{-03}$ |
| 0099503 | Secretory vesicle | 0.55 | $3.00e^{-03}$ |
| 0030054 | Cell junction | 0.41 | $3.00e^{-03}$ |
| 0014069 | Postsynaptic density | 0.77 | $5.00e^{-03}$ |

**Supplemental Figure 1. Validation of the TMT-MS proteomics approach as a quantitative method. A.** Scatter plot of individual patient values for the core AD biomarker qNfL, measured using quantitative assay, and TMT-MS-detected NEFM; (r=0.88, p=1.95e-12). **B.** GO term analysis using STRING was performed on the correlates identified in Figure 2C for each AD core biomarker. The top 3 (ranked by FDR p-value) enriched pathways identified for each core biomarker are listed. **C.** STRING (v11.5) PPI map for qNrgn MS correlates (166 proteins, r ≥ |0.70| and p ≤ 0.01) are shown (medium confidence, disconnected nodes hidden). Proteins of interest are indicated with royal blue arrows: APLP1, amyloid beta precursor like protein 1; NRGN, neurogranin; BIN1, bridging integrator 1; CP, ceruloplasmin. **D.** Top enriched (ranked by FDR) Cellular Component terms for qNrgn MS correlates; relevant terms are shown in blue. **C.** Top-ranked (by FDR) Cellular Component terms for qSyt1 (62 proteins) and qTau (97 proteins) MS correlates are shown (r ≥ |0.70| and p ≤ 0.01). Relevant terms are shown in blue; synapses and synaptic areas are strongly represented.
